# Low generalizability of polygenic scores in African populations due to genetic and environmental diversity

**DOI:** 10.1101/2021.01.12.426453

**Authors:** Lerato Majara, Allan Kalungi, Nastassja Koen, Heather Zar, Dan J. Stein, Eugene Kinyanda, Elizabeth G. Atkinson, Alicia R. Martin

**Affiliations:** Global Initiative for Neuropsychiatric Genetics Education in Research (GINGER), Harvard T.H. Chan School of Public Health, Department of Epidemiology, Boston, MA, USA; MRC Human Genetics Research Unit, Division of Human Genetics, Institute of Infectious Disease and Molecular Medicine, Faculty of Health Sciences, University of Cape Town, Observatory 7925, South Africa; Department of Psychiatry, College of Health Sciences, Makerere University, Kampala, Uganda; Department of Psychiatry, Faculty of Medicine and Health Sciences, Stellenbosch University, Cape Town, South Africa; Mental Health Project, Medical Research Council/Uganda Virus Research Institute (MRC/UVRI) & London School of Hygiene and Tropical Medicine (LSHTM), Uganda Research Unit, Entebbe, Uganda; Department of Psychiatry and Neuroscience Institute, University of Cape Town, Cape Town, South Africa; South African Medical Research Council (SAMRC) Unit on Risk and Resilience in Mental Disorders, Cape Town, South Africa; Department of Paediatrics and Child Health, Red Cross Children’s Hospital and Medical Research Council Unit on Child and Adolescent Health, University of Cape Town, Cape Town, South Africa; Analytic and Translational Genetics Unit, Massachusetts General Hospital, Boston, MA 02114, USA; Stanley Center for Psychiatric Research, Broad Institute of MIT and Harvard, Cambridge, MA 02142, USA; Program in Medical and Population Genetics, Broad Institute of MIT and Harvard, Cambridge, MA 02142, USA

**Keywords:** polygenic scores, Africa, GWAS, health disparities, global health, population genetics

## Abstract

African populations are vastly underrepresented in genetic studies but have the most genetic variation and face wide-ranging environmental exposures globally. Because systematic evaluations of genetic prediction had not yet been conducted in ancestries that span African diversity, we calculated polygenic risk scores (PRS) in simulations across Africa and in empirical data from South Africa, Uganda, and the UK to better understand the generalizability of genetic studies. PRS accuracy improves with ancestry-matched discovery cohorts more than from ancestry-mismatched studies. Within ancestrally and ethnically diverse South Africans, we find that PRS accuracy is low for all traits but varies across groups. Differences in African ancestries contribute more to variability in PRS accuracy than other large cohort differences considered between individuals in the UK versus Uganda. We computed PRS in African ancestry populations using existing European-only versus ancestrally diverse genetic studies; the increased diversity produced the largest accuracy gains for hemoglobin concentration and white blood cell count, reflecting large-effect ancestry-enriched variants in genes known to influence sickle cell anemia and the allergic response, respectively. Differences in PRS accuracy across African ancestries originating from diverse regions are as large as across out-of-Africa continental ancestries, requiring commensurate nuance.

## Introduction

Genome-wide association studies (GWAS) have yielded important biological insights into the heritable basis of many complex traits and diseases (Visscher et al., 2017). However, the vast majority of studies have been conducted in populations of European descent, raising questions about their utility across diverse populations (Manrai et al., 2016; Martin et al., 2019; Morales et al., 2018; Popejoy and Fullerton, 2016; Sirugo et al., 2019). Previous studies have evaluated the generalizability of GWAS by using polygenic risk scores (PRS) to compare the association between genetically predicted versus measured phenotypes in diverse populations.

These studies have found that PRS accuracy decreases with increasing genetic distance between the GWAS discovery and PRS target cohorts (Martin et al., 2017, 2019; Scutari et al., 2016). Since the earliest applications of PRS in human genetics, these concepts--coupled with Eurocentric study biases--have resulted in PRS that are most accurate in European ancestry populations and least accurate in African ancestry populations (International Schizophrenia Consortium et al., 2009). These study biases and phenomena continue to replicate a decade later, with several-fold differences in prediction accuracy of many traits between European and non-European ancestry populations (Martin et al., 2019).

Quantifying PRS generalizability within and among African populations requires considerable nuance as they represent the most genetically diverse populations globally, with more than a million more genetic variants per person than out-of-Africa populations (1000 Genomes Project Consortium et al., 2015). Populations collected even within the same geographic regions of Africa have complex demographic histories with complicated patterns of admixture and population structure (Busby et al., 2016; Choudhury et al., 2020; Pagani et al., 2015; Uren et al., 2016). Further, African ancestry populations experience vastly different environments within versus outside continental Africa as well as more locally among diverse communities, countries, and regions of Africa. These differences provide unique epidemiological opportunities to query the impacts of vastly differing environments on PRS accuracy. Previous empirical analyses and theoretical work fundamentally informs how demographic history and environmental variation interplay to produce PRS heterogeneity in traditionally underserved populations (Mostafavi et al., 2020; de Vlaming et al., 2017; Wang et al., 2020; Wray et al., 2013; Zaidi and Mathieson, 2020).

The inclusion of African ancestry participants in large-scale genetic studies is uniquely important for many reasons. They have the lowest life expectancies globally (Hero et al., 2017; Roser, 2013), receive the lowest access to and quality of medical care in the US (of Health et al., 2017), and are the most underserved by genetic technologies (Martin et al., 2018; Sirugo et al., 2019). A more nuanced understanding of PRS transferability will critically inform which populations are currently the most underserved and thus where building genetic studies and resources will have the biggest benefits globally.

There are also clear benefits to including African populations in statistical genetics efforts. Because humans originated in Africa, populations from Africa have the most genetic diversity among global populations (1000 Genomes Project Consortium et al., 2015; Campbell and Tishkoff, 2008; Henn et al., 2012a), such that more genotype-phenotype associations are expected in Africa than can be found elsewhere. African Americans have been shown to contribute disproportionately to GWAS findings (Morales et al., 2018), making up 2.8% of GWAS participants but contributing 7% of trait associations. African ancestry populations also have shorter blocks of linkage disequilibrium, which improves resolution to fine-map causal variants (Genovese et al., 2010). PRS accuracy is lowest in African ancestry populations due to GWAS study biases (Martin et al., 2019), but when GWAS include these and other diverse populations, PRS predict traits such as schizophrenia more accurately across all populations compared to single-ancestry GWAS (Bigdeli et al., 2019).

In this study, we have investigated how PRS generalize within and among diverse African populations in simulations and with empirical genotype-phenotype data for dozens of quantitative traits. We first simulated causal effects and computed genetic risk prediction accuracy using data from the African Genome Variation Project. We then calculated PRS using publicly available GWAS summary statistics from predominantly European ancestry populations to: 1) quantify PRS accuracy for 5 physical and psychosocial traits among populations in the Drakenstein Child Health Study (DCHS) of South Africa, a birth cohort study; and 2) compare PRS accuracy for 34 quantitative traits across the Ugandan General Population Cohort (GPC) versus ancestrally diverse UK Biobank participants. Our results highlight the disproportionate benefits of genetic studies in diverse African populations to improve trait prediction. Further, while PRS hold promise as biomarkers in precision medicine, a critical prerequisite is equitable accuracy in diverse populations to avoid exacerbating existing health disparities.

## Results

Our study uses both simulation-based and empirical approaches to evaluate the generalizability of PRS across diverse African ancestry populations. An overview of the study design is shown in **Figure 1**, abbreviations are in **Table S1**, and a summary of datasets used in this study are shown in **Table S2**.

**Figure 1.**
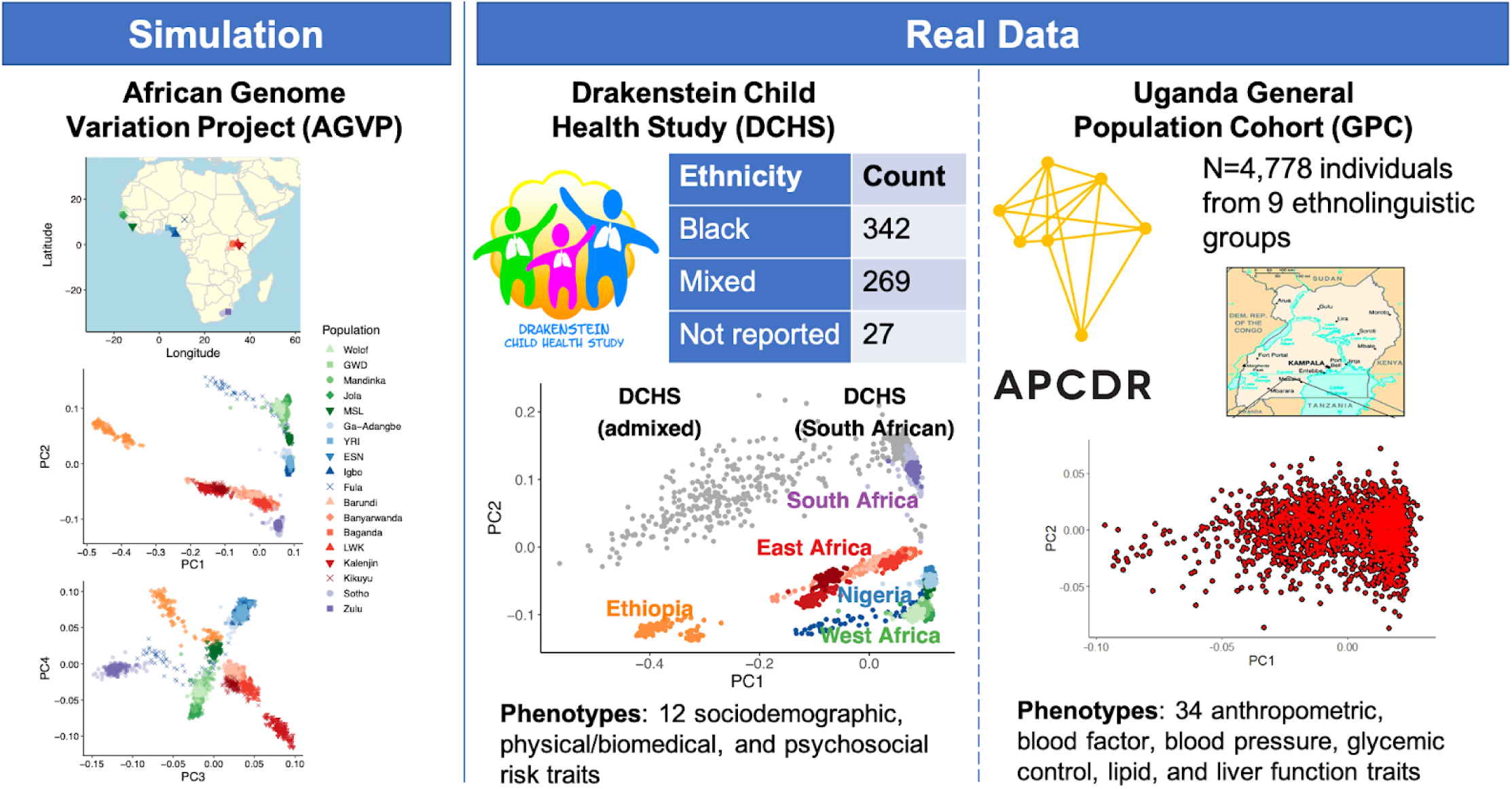
Project overview of genetic and phenotypic datasets used to assess polygenic score generalizability within and across diverse African populations. Using publicly available GWAS data from primarily Eurocentric populations, we measure how polygenic scores perform in Africa. In simulations, we use AGVP genetic data and simulated phenotypes to assess polygenic score generalizability within Africa. In real data, we use two datasets to measure polygenic score accuracy: the South African DCHS cohort data and the Ugandan GPC cohort data.

### Simulated generalizability within and across diverse African populations

We simulated several quantitative traits with varying numbers of causal variants (N = 5; 20; 100; 2,000; 10,000; and 50,000) and heritabilities (h^2^ = 0.1, 0.2, 0.4, and 0.8), then conducted independent GWAS for each scenario in East and West African ancestry populations (**Methods, Figures S1-4**). We calculated the prediction accuracy for PRS derived from the GWAS summary statistics considering ten different p-value thresholds within and across independent target populations from East, West, and South Africa. In general, ancestry-matched results with the sparsest and most heritable genetic architectures produced the highest prediction accuracy. Prediction accuracy was highest with trait h^2^ = 0.8 and fewer than 100 causal variants (**Figure 2A-C**), as indicated by the highest R^2^ and the identification of genome-wide significant associations.

**Figure 2.**
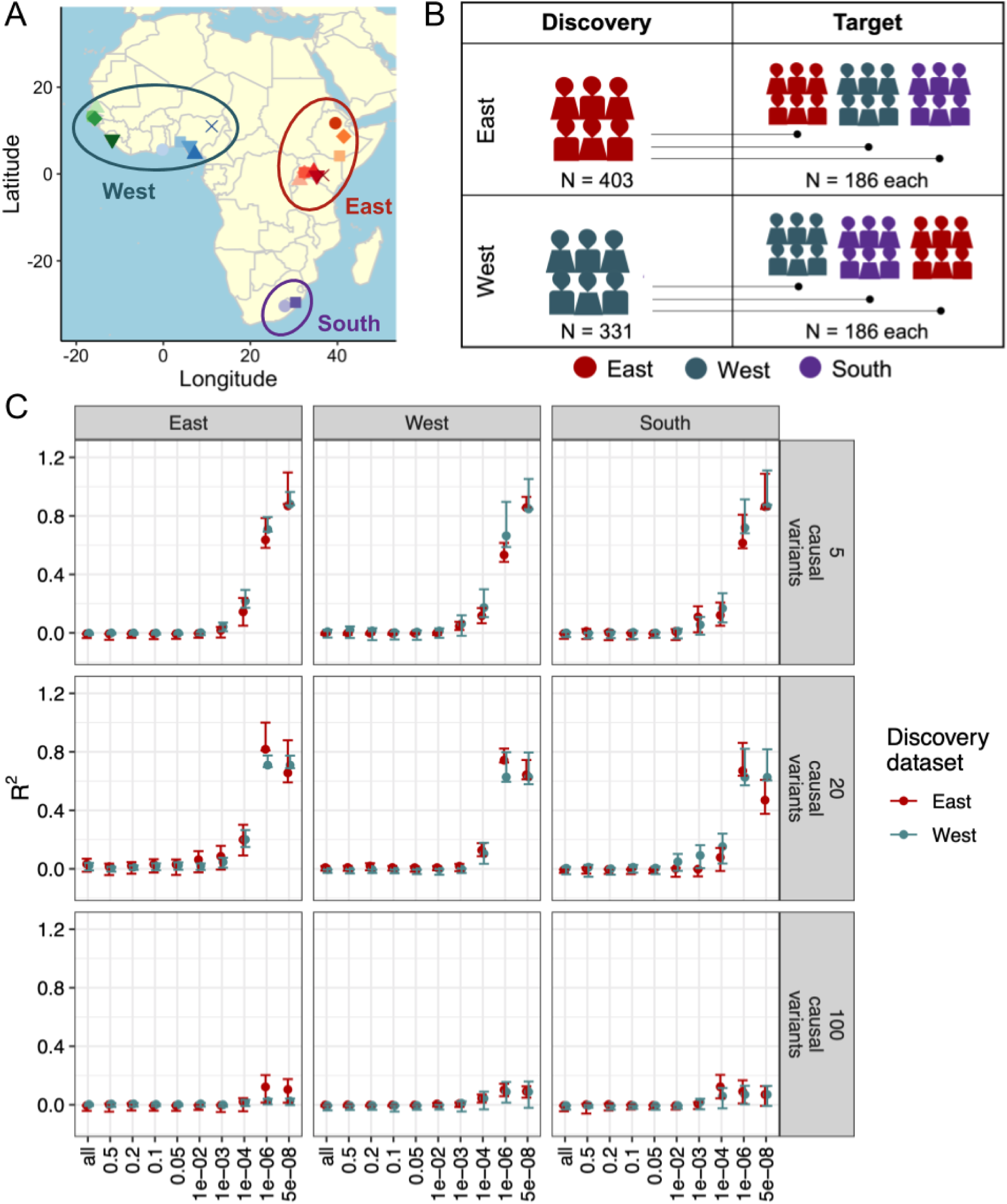
Simulated GWAS and polygenic scores indicate differential prediction accuracy across diverse regions of Africa using genetic data from the AGVP. A) Populations were grouped into East, West, and South based on the United Nations geoscheme groupings. B) GWAS discovery cohorts included East (N = 403) and West (N=331) African individuals, which were independent of each target cohort (N = 186 individuals per region). South Africans were excluded from the discovery population due to the limited total sample size (2 populations and 186 individuals total). C) Predictive accuracy of the simulated quantitative trait at the heritability of 0.8. The predictive accuracy was calculated for six categories of causal variants for the West and East discovery cohorts, across ten *p*-value thresholds.

Conversely, when the number of causal variants exceeded 100, prediction accuracy was negligible (**Figure S4**) because of the small discovery cohort sample sizes, as evidenced by no variants meeting genome-wide significance in these simulations.

Prediction accuracy was highest with 5 and 20 causal variants (**Figure 2C**). The within-ancestry prediction at p-value threshold < 5e-08 and five causal variants were: R^2^ = 0.86, p = 1.74 × 10^−74^ for East discovery - East target scores; R^2^ = 0.85, p = 9.9e-74 for West discovery - West target scores. We observed lower prediction accuracy with ancestry mismatched discovery versus target cohorts at five causal variants and p-value threshold = 1e-6 (R^2^ = 0.66, p = 1.79e-42 for West discovery - West target scores, compared to R^2^ = 0.53, p =1.29e-74 for East discovery - West target scores). The scores in the South target sample were comparable when using East- or West-derived summary statistics (R^2^ = 0.86, p = 5.19e-84 for West-derived summary statistics, and R^2^ = 0.86, p = 1.35e-83 for East-derived summary statistics).

### PRS accuracies in South African populations

While our simulations have shown that PRS generalize poorly across Africa due to substantial genetic diversity and differences across the continent, there is also considerable genetic and environmental diversity within regions and countries. We quantified PRS accuracy for a range of measured phenotypes in mothers genotyped in the DCHS cohort in South Africa, including several sociodemographic, physical/biomedical, and psychosocial risk traits (**Table S3**). The DCHS cohort consists of participants with multiple ancestry groups that include an admixed population with ancestry from multiple continents as well as a population with almost exclusively African population. These ancestry groups correlate with self-reported “Mixed” and “Black/African” ethnicities, respectively (**Figure S5**). We computed PRS for maternal height, depression, psychological distress, alcohol consumption, and smoking in DCHS overall, by ethnic group, and by ancestry within the Mixed ethnic group (**Methods**).

Across all genetically predicted phenotypes, only height was significantly predicted (**Figure S6**). We predicted height more accurately in the Mixed versus Black/African ethnic groups (R^2^ = 0.099, 95% bootstrapped CI = [0.012, 0.18], p =1.5e-7 versus R^2^ = 0.021, 95% CI = [-0.031, 0.043], p = 5.27e-3, respectively). We also expect that PRS accuracy increases with decreasing African ancestry within the Mixed ethnic group as has been shown previously in admixed African populations (Bitarello and Mathieson, 2020); we find suggestive evidence consistent with this trend when partitioning the Mixed group into two bins along PC1 (R^2^ = 0.091, 95% CI = [-0.04, 0.17], p = 6.4e-4 in lower half of PC1 with more African ancestry vs R^2^ = 0.12, 95% CI = [-9.0e-4, 0.21], p=5.7e-5 with more out-of-Africa ancestry), although small sample sizes limit definitive comparisons (N = 137 in each PC1 bin). Our results are consistent with variable prediction accuracy among diverse African ancestry groups within South Africa and insignificant prediction in African populations for all but the most heritable and accurately predicted traits elsewhere.

### Variable phenotypic and genetic similarities across the Uganda General Population Cohort (GPC) and UK Biobank

#### Lower phenotypic correlations in Uganda GPC suggest higher contributing environmental effects

We next investigated phenotypic similarities within and across the Uganda GPC and UK Biobank participants because these are two of the largest cohorts with dozens of traits measured in African ancestry individuals. We first considered overall cohort differences between these cohorts--the Uganda GPC enrolled participants using a house-to-house study design and generated genetic data on 5,000 adults from rural villages in southwestern Uganda (Asiki et al., 2013), while the UK Biobank enrolled 500,000 people aged between 40-69 years in 2006-2010 from across the country (**Methods** (Bycroft et al., 2018)). Previous studies have reported higher rates of infectious diseases (e.g. HIV, hepatitis B and C) in the Uganda GPC than would be expected in the UK Biobank (Asiki et al., 2013). There are many additional potential environmental explanations for mean shifts in phenotypes, such as dietary, food security, and age differences contributing to considerable BMI differences across cohorts (μ = 21.3 and σ = 3.8 in Uganda GPC versus μ = 27.4 and σ = 4.8 in UK Biobank, p < 2.2e-16). To quantify comparisons while controlling for demographic differences for each of the 34 quantitative traits measured in both cohorts, we first mean centered each phenotype and regressed out the effects of age and sex within each cohort. Next, we then compared the distributions and variances of each phenotype across cohorts via Kolmogorov-Smirnov and F-tests, respectively (**Table S4**). Given the large sample sizes, all K-S tests were significantly different, with several phenotypes showing distributional and variance differences of considerable magnitude (**Figure S7** and **Table S4**, e.g. Bilirubin, BASO, HbA1c, ALP, EOS, TG, and NEU).

We next analyzed how similar the relationships are between phenotypes across datasets. Similar trends emerge overall, with distances across variance-covariance matrices for these cohorts showing evidence of significant correlation (Mantel test Z-statistic = 0.73, p < 1e-4).The correlations among phenotypes are slightly higher overall in the Uganda GPC than in UK Biobank, both among related and unrelated individuals, as expected from a household versus volunteer-based design (**Figure 3B, Figure S8**). More specifically, we see consistent correlations among combinations of phenotypes including SBP and DBP; RBC, Hb, and HCT; Cholesterol and LDL; WC, BMI, WT, and HC; MCHC, MCH, and MCV; GGT, ALT, AST, and ALP; and MONO, NEU, and WBC with high overall correlations across these datasets for these traits (**Figure 3A-B**, see abbreviations in **Table S1**). Some pairs of traits, however, have significantly different correlations across datasets. The largest difference in phenotypic correlations across datasets is between ALP and WT (ρ = 0.11, p < 2.2e-16 in UK Biobank versus ρ = −0.36, p < 2.2e-16 in Uganda GPC).

**Figure 3.**
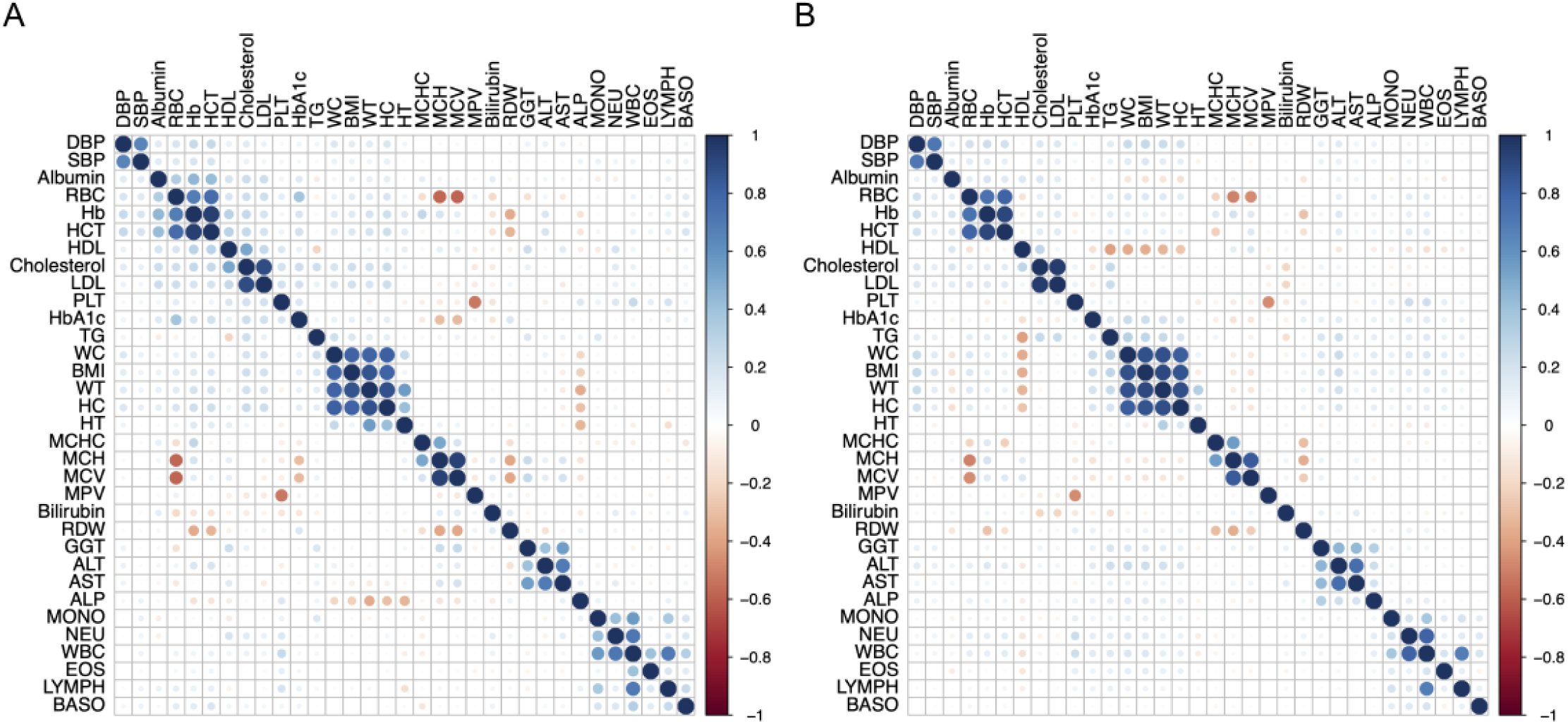
Phenotype and genotype correlations among 33 quantitative traits measured in the Uganda GPC data and the UK Biobank. A) Phenotypic correlations measured in traits in the Uganda GPC among unrelated individuals. B) Phenotypic correlations in the UK Biobank European ancestry unrelated individuals. A-B) Phenotypes were mean centered and adjusted for age and sex within each cohort prior to correlation analysis. The order of each phenotype correlation is determined by hierarchical clustering in the Uganda GPC.

Our next goal was to compare trait heritability estimates in the UK Biobank versus Uganda GPC data (**Methods**). However, the sample size and study design differences between these cohorts required the application of different methods that limit comparability. Specifically, the household design of Uganda GPC included smaller sample sizes with more relatives in which family-based heritability estimates are most appropriate, whereas the large sample size and volunteer design in UK Biobank makes SNP-based heritability estimates from unrelated individuals most appropriate. **Figure S9** compares heritability estimates across traits in the UK Biobank versus Uganda GPC using these approaches (Gurdasani et al., 2019). As expected from the differences in the methods, study designs, and sample sizes, we find higher but noisier estimates in Uganda GPC for most traits, consistent with expectation from family-based versus unrelated heritability estimates across these two studies.

### African genetic risk predictions from European ancestry GWAS data are remarkably inaccurate

To understand baseline trans-ethnic PRS accuracy using a typical approach, we predicted 32 traits in the Uganda GPC using GWAS summary statistics from the UK Biobank European ancestry individuals. While several traits were significantly predicted across ancestries, prediction accuracy was low for most traits (**Figure S10**); the most accurate PRS was for MPV, (R^2^ = 0.036, 95% CI = [0.0069, 0.063], p = 5.73e-7) while the average variance explained across all traits was less than 1% (mean R^2^ = 0.007). To assess the relative effects of ancestry versus cohort differences on decreases in prediction accuracy across populations, we next withheld 10,000 European ancestry individuals from UK Biobank for use as a target cohort, reran all GWAS, then used individuals with diverse continental ancestries in the UK Biobank as target populations (EUR = Europeans withheld from the GWAS, AMR = admixed American, MID = Middle Eastern, CSA = Central/South Asian, EAS = East Asian, and AFR = African, **Figure S11**), subcontinental African ancestries in the UK Biobank (Ethiopian, Admixed, South, East, West African ancestries, **Figure S12**), as well as the Uganda GPC (**Figure 4A, Table S4**).

**Figure 4.**
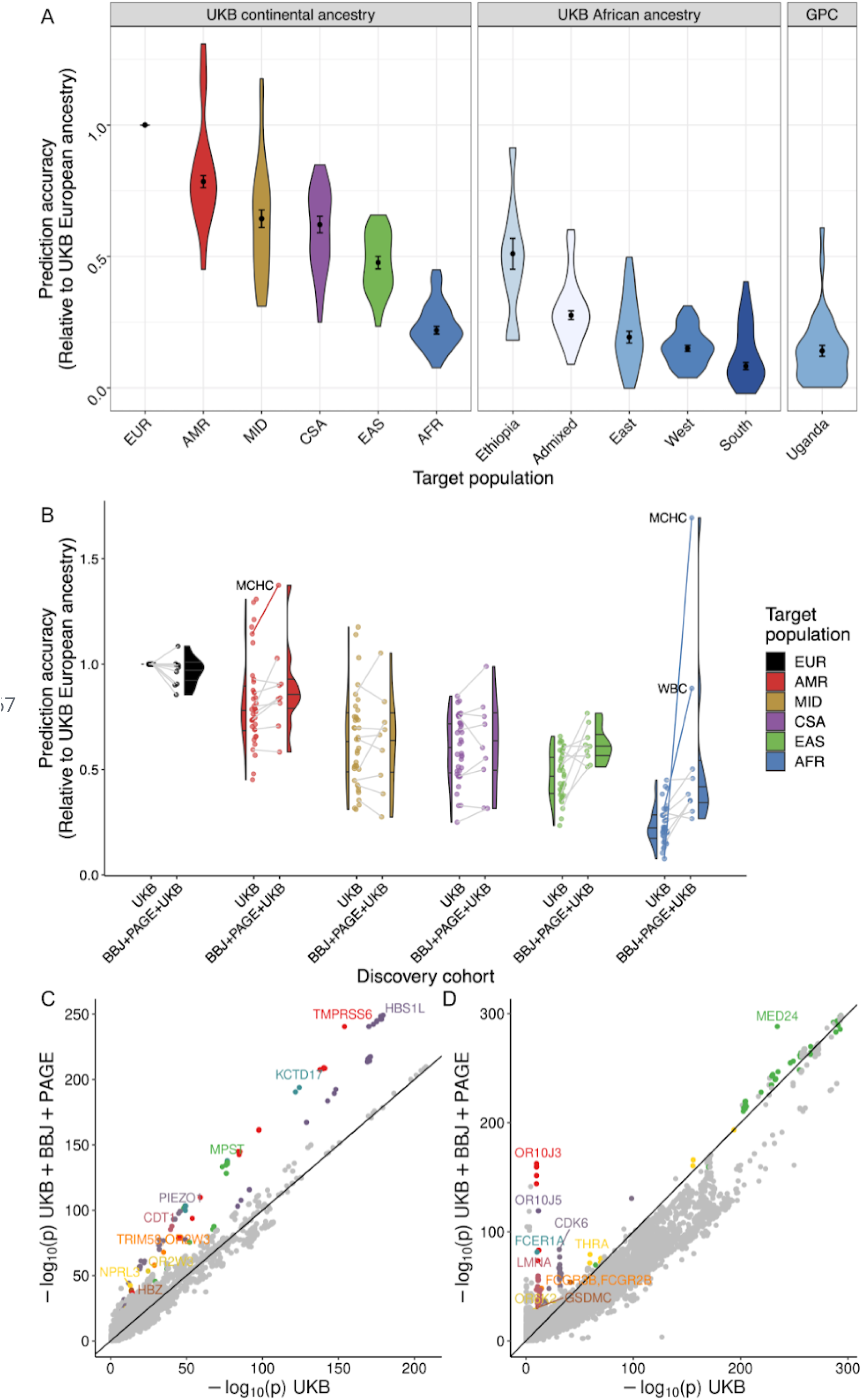
PRS accuracy and corresponding genetic variant contributions for up to 34 traits within and across diverse ancestries. A) PRS accuracy relative to European ancestry individuals in diverse target ancestries. Discovery data consisted of GWAS summary statistics from UK Biobank (UKB) European ancestry data. Target data consisted of globally diverse continental ancestries (including withheld European target individuals) and regional African ancestry participants from UKB, or unrelated individuals from the Uganda GPC cohort. Traits were filtered to those with a 95% confidence interval range in PRS accuracy < 0.08. B) PRS accuracy from a homogeneous versus multi-ancestry discovery dataset. GWAS discovery data consisted of summary statistics from UKB European ancestry data only or from the meta-analysis of UKB, BioBank Japan (BBJ), and Population Architecture using Genomics and Epidemiology (PAGE). Target populations are from the UKB. Lines connect the 10 traits available in both discovery cohorts to indicate how accuracy changed for the same trait in the UKB only versus meta-analyzed discovery data, while half violin plots show the distribution across all phenotypes in each discovery cohort. When lines are missing, the trait is absent in PAGE. Trait outliers are labeled in text and with solid lines. A-B) Relative PRS accuracies are compared to the maximum for each trait in target samples withheld from discovery consisting of UKB European ancestry individuals. To simplify comparisons, only the polygenic scores with the highest prediction accuracy are shown here. Colors in these two panels correspond to the same continental ancestries. C-D) Trait-specific genetic outlier plots. QQ-like plot showing p-values in UKB only versus multi-cohort meta-analysis of UKB, BBJ, and PAGE. The ten regions that are genome-wide significant in both dataset and show the most significant differences are colored and labeled for: C) MCHC, and D) WBC.

Among continental ancestries, we computed R^2^ and 95% confidence intervals for each trait (**Figure S13**), then computed median relative accuracy (RA) compared to Europeans and median absolute deviation (MAD) across all traits. We predict these traits most accurately in EUR (RA = 1, MAD = 0), followed by AMR (RA = 0.784, MAD = 0.023), MID (RA = 0.643, MAD = 0.034), CSA (RA = 0.621, MAD = 0.031), EAS (RA = 0.477, MAD = 0.024), and AFR (RA = 0.219, MAD = 0.014) (**Figure 4A**). We next compared prediction accuracy within African ancestry populations. Because some PRS accuracy estimates were noisy due to small sample sizes in UK Biobank Africans (especially Ethiopian and South African ancestry individuals, **Table S4**), we restricted analyses to those traits predicted with a 95% confidence interval < 0.08. Among these traits, we predicted most accurately those with Ethiopian ancestry (RA = 0.511, MAD = 0.059), followed by recently admixed individuals with West African and European ancestry (RA = 0.276, MAD = 0.016), East African ancestry (RA = 0.193, MAD = 0.023), West African ancestry (RA = 0.150, MAD = 0.012), and South African ancestry (RA = 0.083, MAD = 0.014) (**Figure 4A**). These results track with genetic distance and population history; the highest prediction accuracy identified in Ethiopians is expected given closer genetic proximity to European populations relative to other Africans due to back-to-Africa migrations influencing population structure there (Henn et al., 2012b; Hodgson et al., 2014; Pagani et al., 2015). The lowest prediction accuracy is in populations with southern African ancestry, consistent also with higher genetic divergence from European populations and more genetic diversity overall (Busby et al., 2016; Choudhury et al., 2020; Henn et al., 2011).

#### Lower prediction accuracy across ancestries than across cohorts

To compare prediction accuracy among similar ancestry participants from different cohorts, we next computed PRS for 32 traits using GWAS summary statistics from UK Biobank Europeans in two target populations: UK Biobank participants with East African ancestry versus Uganda GPC. As expected, prediction accuracy in these populations is very low across all traits in both cohorts and only slightly higher in the UK East African ancestry individuals than in the Uganda GPC individuals (mean R^2^ = 0.017, sd = 0.013 versus mean R^2^ = 0.012, sd = 0.010, respectively, **Figure S14**). Across traits, the differences in PRS accuracy across cohorts but within the same ancestry are much smaller than the differences across ancestries but within the UK Biobank, indicating that ancestry has a larger impact on genetic risk prediction than cross-cohort differences analyzed here. Smaller effects on genetic prediction accuracy differences across cohorts may be attributable to environmental differences, such as higher rates of malnutrition and infectious diseases previously reported in Uganda and in the GPC (Asiki et al., 2013; Nalwanga et al., 2020).

#### Improved African genetic risk prediction accuracy with multi-ethnic GWAS summary statistics

We next maintained the target populations but varied the discovery cohort to determine how more diverse GWAS impacts PRS accuracy for these phenotypes in diverse populations. Specifically, we computed PRS accuracy in diverse target populations in the UK Biobank (**Table S5**) using one of two discovery cohorts: the UKB European-only cohort versus diverse discovery cohorts combined via meta-analysis (**Table S6**). Meta-analyzed GWAS summary statistics come from several cohorts, including the UK Biobank (UKB), Biobank Japan (BBJ) (Nagai et al., 2017), Population Architecture Using Genomics and Epidemiology (PAGE) Consortium (Wojcik et al., 2019), and Uganda Genome Resource (UGR) (Gurdasani et al., 2019). For each trait, discovery cohort, and target cohort combination, we normalized the PRS R^2^ values from the p-value threshold that explained the maximum phenotypic variance with respect to the prediction accuracy in the European target cohort using UK Biobank summary statistics only, then computed relative accuracies as before.

We find that prediction accuracy improves the most across populations when using a discovery cohort consisting of GWAS summary statistics meta-analyzed across the UKB, BBJ, and PAGE cohorts (**Figure 4B**), but not the UGR data (**Figure S15**). Instead, meta-analyzing the UGR data with UKB did not improve prediction accuracy for any population and most notably decreased accuracy in African ancestry target populations (discovery UKB median RA = 0.22, UGR+UKB median RA = 0.15, **Figure S16**). We hypothesize that this can be explained by the relatively small sample size of UGR adding more noise than signal compared to the other relatively large discovery datasets, but another explanation could come from environmental heterogeneity. When predicting traits using the UKB, BBJ, and PAGE meta-analysis as a discovery cohort, we find that prediction accuracy increases most for the AMR, EAS, and AFR target populations, which more closely resemble the ancestry patterns of PAGE and BBJ (**Figure 4B**). These findings are consistent with ancestry-matched discovery data disproportionately improving prediction accuracy in the corresponding target population (Bigdeli et al., 2019; Lam et al., 2019; Martin et al., 2019).

#### Large-effect population-enriched genetic variants drive heterogeneity in polygenic score accuracy for blood panel traits

We find that PRS accuracy improvements from higher diversity in the discovery cohorts vary across traits, with the largest increases seen in MCHC and WBC. We searched for specific genetic loci that could explain this pattern by comparing the significance of genetic associations in UKB alone versus the meta-analysis of UKB, BBJ, and PAGE (**Table S6**). For MCHC and WBC in particular, the genetic variants contributing to these improved PRS consist of several well-known population-enriched variants (**Figure 4C** and **4D**). For example, genetic variants that disproportionately explain population-specific risk for MCHC include variants previously associated with hemoglobin concentration, including rs9399137 upstream of *HBS1L* and *MYB* in a study of sickle cell anemia (p = 5.24e-249 and β = 0.0783 in the meta-analysis) (Lettre et al., 2008), rs855791 in *TMPRSS6* (p = 3.49e-241, β = 0.0692) (Benyamin et al., 2009; Chambers et al., 2009), and rs551118 upstream of *PIEZO1* and *CDT1* (p = 5.18e-100, β = −0.0451) (Astle et al., 2016) (**Table S7**). Associations with WBC tend to show more population-enriched associations as shown in the meta-analysis (**Figure 4D**), including rs3936197 in *MED24* (p = 5.18e-289, β = −0.0772), rs58650325 near the high affinity IgE receptor *FCER1A* that initiates the allergic response (1.57e-163, β = −0.097, also close to *OR10J3*), and rs11533993 in *CDK6* (p = 1.55e-84, β = −0.0799). Thus, genetic architecture and population genetic considerations are important to bear in mind when considering the generalizability of polygenic scores.

## Discussion

PRS have been proposed as genetic biomarkers for use in preventative medicine (Khera et al., 2018; Knowles and Ashley, 2018), but are currently limited by low accuracy across populations especially in African ancestry populations (Martin et al., 2019; Sirugo et al., 2019). This study has enabled unique insights into PRS transferability within and among diverse continental African populations as well as among African ancestry populations living in considerably different environments. We demonstrate looming challenges for applying current PRS in African ancestry populations; because relatively few genetic studies have been conducted in African populations coupled with their uniquely deep population histories, PRS accuracy is low but widely variable. Differences in PRS accuracy across diverse African ancestries from different regions can be larger than across out-of-Africa continents. This is particularly problematic as widely-used algorithms that guide health decisions already have ingrained racial biases (Obermeyer et al., 2019), warning of compounding challenges with implementation. We demonstrate that there are clear steps the field can take to work against these biases. Specifically, including ancestrally diverse populations in GWAS discovery cohorts improves accuracy for all populations and especially underrepresented populations more than conducting similarly sized studies with only European ancestry cohorts.

Another advantage of using GWAS from globally diverse populations to compute PRS is the routine inclusion of population-enriched variants. Clear examples such as African-enriched variants in *APOL1* and *G6PD* have been shown to contribute especially high risk of chronic kidney disease and to missed diabetes diagnosis, respectively (E et al., 2018; Rotimi et al., 2017). These examples highlight the importance of studying diverse populations to predict genetic risk of disease equitably by aggregating variants across the spectrum of allele frequencies and effect sizes in different populations. Relevant to the traits studied in genetic analyses here, hematological differences such as anemia are more common in lower income countries in Africa and in African ancestry populations elsewhere compared to European ancestry populations in high income countries, particularly among older individuals. These hematological differences potentially arise in part due to genetic variation as well as the higher prevalence of infectious diseases and pathogens, poorer nutritional status, and altitude (Mugisha et al., 2013, 2016). Here, we show that variants influencing risk of beta thalassemia disproportionately increase PRS accuracy for hemoglobin variation particularly in African ancestry populations. The inclusion of population-enriched variants in PRS could eliminate genetic justifications for race-based medicine, which problematically reinforces implicit racial biases by overemphasizing the link between genetics and race despite the fact that there is more genetic variation within than between populations (Cerdeña et al., 2020).

In addition to reduced PRS accuracy with ancestral distance from GWAS cohorts, genetic nurture, social genetic, and environmental effects can also contribute to low portability of PRS across populations (He et al., 2019; Mostafavi et al., 2020), with some interventions modulating health along PRS strata (Barcellos et al., 2018). In this study, however, ancestry appears to have a larger effect on portability than cohort differences overall. An important distinction when comparing the magnitude of these and other non-genetic effects in other studies is that the traits most accurately genetically predicted here were primarily anthropometric and blood panel traits. When analyzing traits with more sociodemographic influences in increasingly diverse populations, population stratification, confounding, and study design considerations are thornier issues (Kerminen et al., 2019; Novembre and Barton, 2018; Zaidi and Mathieson, 2020). PRS accuracy comparisons across ancestrally similar but environmentally diverse populations are especially important for medically actionable traits. For example, particularly low PRS portability for triglycerides (TG) from European to the Uganda GPC resulted at least in part from effect size heterogeneity that has previously been connected to pleiotropic and gene * environment effects; specifically, most non-transferable genome-wide significant associations with TG showed pleiotropic associations with BMI in Europeans but not Ugandans (Kuchenbaecker et al., 2019).

While PRS currently have limited portability, increased diversity in genetic studies is already decreasing prediction accuracy gaps across populations (Bigdeli et al., 2019; Kuchenbaecker et al., 2019). This is consistent with causal genetic effects tending to be similar across populations but with LD and allele frequency differences modifying marginal effect size estimates (Martin et al., 2019). This is also consistent with trans-ethnic genetic correlations tending to be close to or not significantly different from 1 (Brown et al., 2016; Shi et al., 2020). The most rapid path to closing gaps in PRS transferability is to increase the inclusion of GWAS participants from populations most divergent from those already routinely studied. As empirically demonstrated here, when comparing PRS accuracy calculated from diverse cohort meta-analysis versus data from Europeans only, large-scale GWAS with diverse African populations will most rapidly reduce portability gaps across global populations because they have the most genetic diversity, most rapid linkage disequilibrium decay, and highest genetic divergence from the best studied populations. Major efforts underway such as the Human Hereditary and Health in Africa Initiative, PAGE, All of Us, and NeuroGAP programs (All of Us Research Program Investigators et al., 2019; Hindorff et al., 2018; Mulder et al., 2018; Stevenson et al., 2019; Wojcik et al., 2019) are especially promising for rectifying current PRS gaps and missed scientific opportunities by increasing inclusion of diverse African participants.

Beyond expanding on diversity by increasing the number of study participants in large-scale studies, it is equally important to diversify researchers working on genomics studies. Currently, the vast majority of researchers in genomics studies are of European ancestry (Ginther et al., 2011; Hamrick, 2019; Hoppe et al., 2019), paralleling the over-representation of European-ancestry individuals in genomic studies. The exclusion of African researchers leads to the disparity in research leadership and reduced scientific output from African researchers (Bentley et al., 2020). Efforts such as the NeuroGAP Global Initiative for Neuropsychiatric Genetics Education and Research (GINGER) program (van der Merwe et al., 2018), which provides mentorship and training for early-career investigators on the African continent (particularly in Uganda, Kenya, Ethiopia and South Africa, including several of this study’s authors), are important in moving toward a more inclusive and representative research community.

## Conclusion

Previous studies that have examined PRS accuracy across globally diverse ancestry groups have demonstrated that accuracy is lowest in African ancestry samples. However, the extent to which this accuracy varies within African-ancestry populations has not been previously investigated. Our findings that prediction accuracy varies by African-ancestry populations is a clear reflection of the vast genetic diversity of the continent. It is therefore critically important to create well-powered GWAS that reflect the full range of diversity within Africa.

## Materials and Methods

### Genetic and Phenotypic Data

Total counts of individuals by population and/or study are shown in **Table S2**.

#### 1000 Genomes Project

1000 Genomes Project data from the phase 3 integrated call set was accessed and used as a reference panel and for phasing and imputation. (1000 Genomes Project Consortium et al., 2015)

#### Human Genome Diversity Project (HGDP)

Genotype data for samples from HGDP was publicly available on the Illumina HumanHap650K GWAS array on hg18 (Li et al., 2008). We lifted over the genotype data to the hg19 genome build using hail (http://hail.is).

#### African Genome Variation Project (AGVP)

As described previously (Gurdasani et al., 2015), the AGVP data consists of dense genotype data from 1,481 individuals from 18 ethno-linguistic groups from Eastern, Western, and Southern Africa when including the Luhya and Yoruba from the 1000 Genomes Project (1000 Genomes Project Consortium et al., 2015). When accessed from the European Genome-Phenome Archive (EGA), “Ethiopian” is the provided population label encompassing the Oromo, Amhara, and Somali groups. After collapsing these groups and counting the 1000 Genomes data separately, 1,307 individuals from 14 populations are uniquely represented in AGVP, and 2,504 individuals from 26 populations are represented in the 1000 Genomes Project data (661 individuals from 7 populations are in the AFR super population grouping).

#### Drakenstein Children’s Health Study (DCHS) in South Africa

The DCHS is an ongoing, multidisciplinary population-based birth cohort study in the Drakenstein area in Paarl (outside Cape Town, South Africa) (Stein et al., 2015; Zar et al., 2015, 2019). After providing informed consent, pregnant women were enrolled during their second trimester (20–28 weeks gestation); maternal-child dyads were then followed through childbirth and longitudinally thereafter. Enrollment occurred from March 2012 to March 2015 at two primary health care clinics - TC Newman (serving a predominantly mixed ancestry population) and Mbekweni (serving a predominantly Black African population). Women were eligible to participate in the DCHS if they attended one of the study clinics, were at least 18 years of age and intended to remain residing in the study area.

#### Uganda General Population Cohort (GPC)

The rural Uganda GPC of MRC/UVRI & LSHTM Uganda Research Unit was set up in 1989 initially to monitor the HIV epidemic among adults, children, and adolescents, but its mandate has since expanded to include other medical conditions (Asiki et al., 2013). The ‘original GPC’ is located in the sub-county of Kyamulibwa in rural south-western Uganda with activities having recently been expanded to the neighbouring two peri-urban townships of Lwabenge and Lukaya. The ‘original GPC’ includes about 10,000 adults and about 10,000 children and adolescents. In 2011, genotype data was generated on more than 5,000 adult participants from nine ethnolinguistic groups using the Illumina HumanOmni2.5 BeadChip at the Sanger Wellcome Trust Institute (Asiki et al., 2013; Heckerman et al., 2016).

#### UK Biobank (UKB)

The UK Biobank enrolled 500,000 people aged between 40-69 years in 2006-2010 from across the country, as described previously (Bycroft et al., 2018). A more detailed description of the cohort is available on their website: https://www.ukbiobank.ac.uk/. We analyzed phenotypes that overlapped with those studied in the Uganda GPC.

### Ancestry analysis in the UK Biobank

As described previously (Bycroft et al., 2018), the UK Biobank consists of approximately 500,000 participants of primarily European ancestry who have thousands of measured or reported phenotypes. To assess polygenic score accuracy across diverse ancestries, we identified populations of ancestral groups at two levels: 1) among continental groups, and 2) among regions in Africa. To define continental ancestries, we first combined reference data from the 1000 Genomes Project and HGDP. We combined these reference datasets into continental ancestries according to their corresponding meta-data (**Table S5**). We then ran PCA on unrelated individuals from the reference dataset. To partition individuals in the UK Biobank based on their continental ancestry, we used the PC loadings from the reference dataset to project UK Biobank individuals into the same PC space. We trained a random forest classifier given continental ancestry meta-data (AFR = African, AMR = admixed American, CSA = Central/South Asian, EAS = East Asian, EUR = European, and MID = Middle Eastern) based on the top 6 PCs from the reference training data. We applied this random forest to the projected UK Biobank PCA data and assigned initial ancestries if the random forest probability was >50% (similar results obtained for p > 0.9), otherwise individuals were dropped from further analysis.

We next further partitioned African ancestry individuals using the same random forest approach as above but without further probability thresholding using African ancestry reference data from AGVP, HGDP, and the 1000 Genomes Project. We partitioned these reference data into UN regional codes with an additional region for Ethiopian populations given their unique population history and collapsing in African Genome Variation Project data (Admixed, Central, East, Ethiopia, South, and West Africa), as shown in **Table S5**. PCA with reference data at the continental and subcontinental level within Africa are shown in **Figures S10-11**.

### Phasing and imputation

We used the Ricopili pipeline to conduct pre-imputation QC and perform phasing and imputation for AGVP and the Uganda GPC (Lam et al., 2020). This pipeline was also used on the DCHS data, as described previously (Duncan et al., 2018). Briefly, we phased the data using Eagle 2.3.5 and imputed variants using minimac3 in chunks ≥ 3 Mb. The 1000 Genomes phase 3 haplotypes were used as the reference panel for phasing and imputation. For the AGVP, we used strict best guess genotypes where a variant was called if it had a probability of *p* > 0.8 and a missing rate less than 0.01 and MAF > 5%. Then, variants with MAF < 0.001 were excluded from the dataset. For Uganda GPC, we used combined best guess genotypes where a variant was called if it had a probability *p* > 0.8 or set to missing otherwise. Then, SNPs were filtered to keep sites with missingness < 0.01 and MAF > 0.05. We used genotype dosages when computed PRS.

### PCA

Only SNPs with high imputation quality (INFO>0.8) were considered for principal component analysis. We computed the first 20 principal components using plink with the --pca flag for autosomal SNPs MAF > 0.05 and individual missingness < 0.05.

### Simulation setup

To test the PRS prediction accuracy within and across African populations, we simulated four quantitative traits while varying heritabilities (*h*^*2*^ = 0.1, 0.2, 0.4 and 0.8) as follows:

We randomly assigned an effect size to 5, 20, 100, 2,000, 10,000 and 50,000 causal variants, respectively. The causal effect was calculated based on the relationship between effect size and minor allele frequency as shown by (Schoech et al., 2019). We then calculated an individual’s ‘true’ polygenic risk as the sum of all causal effects using the --score flag in PLINK v1.07B (Chang et al., 2015). True polygenic scores were standardized to a mean of zero and standard deviation of 1. To account for the contribution of environmental risk factors, we assigned environmental effects from a normal random distribution (mean = 0 and sd = 1). The phenotype was generated according to its heritability as the weighted sum of the true polygenic risk and a random environmental effect as below:

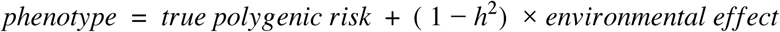

We then conducted GWAS for the simulated phenotype by splitting the AGVP dataset into three groups broadly representing the three geographical areas where samples were obtained from: East (n = 589), West (n = 517) and South Africa (n = 186, **Figure 2A**). To allow for the quantification of PRS prediction accuracy across the geographical regions, each group was further split into discovery and target cohorts. The size of the target cohorts was maintained at n = 186 across all groups, while the discovery cohort consisted of all remaining individuals (East n = 403, West n = 331, and no South Africans). We conducted a linear regression for all the simulated traits for the East and West discovery datasets, controlling for the first 20 principal components.

In PLINK v1.07, independent SNP sets were obtained for each discovery cohort by clumping SNPs from corresponding summary statistics files with an R^2^ value greater than 0.1 using in-sample LD and within 500 kb of each other. The effect sizes from the SNP set was used as weights to compute PRS for all three of our target datasets for a range of *P*-values (5e-08, 1e-06, 1e-04, 1e-03, 1e-02, 0.05, 0.1, 0.2, 0.5 and all). PRS was calculated as the sum of all SNPs multiplied by their effect sizes. We calculated PRS for each of the target datasets using the summary statistics from the discovery dataset GWAS (**Figure 2B**).

### Heritability estimation

For the Ugandan GPC, we relied on heritability estimates of 34 quantitative traits computed previously (Gurdasani et al., 2019). For UK Biobank, we computed heritability estimates for the same traits using LD score regression with the default model (i.e. without any functional annotations) (Bulik-Sullivan et al., 2015) and using used population-matched LD score references from European populations downloaded from the authors’ website (https://data.broadinstitute.org/alkesgroup/LDSCORE/).

### Polygenic score calculation

All PRS were calculated using a pruning and thresholding approach implemented either in plink2 or in hail using custom scripts. All clumping was done in plink2 using an LD threshold of *r*^2^ = 0.1 and a window size of 500 kb with discovery cohort population-specific reference panels. We calculated PRS using plink2 with the --score and --q-score-range flags for AGVP simulations and DCHS. We wrote custom scripts in hail (http://hail.is) to calculate PRS in the Uganda GPC and UK Biobank data due to the larger sample sizes (see **Web resources**). For imputed genotypes, we used SNP dosages in PRS calculations. We computed 10 PRS for each analysis using the following p-value thresholds: 1, 0.5, 0.2, 0.1, 0.05, 0.01, 1e-3, 1e-4, 1e-6, 5e-8. The PRS that explained the most phenotypic variance is shown in most figures.

We calculated PRS accuracy for continuous traits computed with custom scripts in R (**Web resources**). For AGVP simulations and DCHS (because all participants were mothers of a similar age), we included the first 10 PCs as covariates when computing the partial R^2^ specifically attributable to the PRS. For Uganda GPC data, we included age, sex, and the first 10 PCs when computing partial R^2^ of the PRS. For consistency with the GWAS that were run in UK Biobank previously (Howrigan, 2017) and here with a holdout target set, we included, age, sex, age^2^, age*sex, age^2^*sex, and the first 10 PCs as covariates when computing the PRS partial R^2^. (The UKB European GWAS included 20 PCs, but fewer were used here due to the particularly small sample sizes of some other target ancestry groups, **Table S5**, coupled with minimal population structure observed in PCs lower than PC10).

#### Meta-analysis

We used plink2 to conduct inverse variance-weighted meta-analysis across GWAS summary statistics with the --meta-analysis option.

#### LD reference panels and clumping

All PRS calculations required an LD panel for clumping. Our analyses used in-sample LD where feasible and reference panel data as a proxy with ancestry matching from the 1000 Genomes Project phase 3 data when individual-level data was unavailable. We weighted the ancestral representation of each population per trait matching at the continental level. We matched individuals as follows:

| Cohort | 1000 Genomes phase 3 reference data |
| --- | --- |
| BBJ | East Asian (EAS) |
| UKB | European (EUR) |
| UGR | African (AFR) |
| PAGE | Proportional weighting of AFR, EAS, AMR (depending on trait, see **Table S6** description for more detail) |

We then used the maximal number of individuals available when weighting proportionally to construct this reference panel. For example, in the meta-analysis of height across the UKB, BBJ, and PAGE cohorts, UKB has the largest sample size in the discovery cohort (N = 350,353), so all Europeans from 1000 Genomes were included in the reference panel (N = 503), then a random sampling of EAS, AFR, and AMR individuals were included proportionally to the overall diversity of the discovery cohorts in the meta-analysis.

## Supporting information

Supplementary Figures 1-16, captions for Tables 1-7

## Acknowledgements

We thank Lori Chibnik, Bizu Gelaye, Kristi Post, and Courtney White for facilitating the GINGER program and making this work possible. This work was supported by funding from the National Institutes of Health (K99MH117229 to A.R.M.; K01MH121659 and T32MH017119 to E.G.A.). UK Biobank analyses were conducted via application 31063. The DCHS is funded by the Bill and Melinda Gates Foundation (OPP 1017641). Additional support for HJZ and DJS, and for the research reported in this publication, was provided by the South African Medical Research Council (SAMRC). The SAMRC provided additional support through its Division of Research Capacity Development under the National Health Scholarship programme from funding received from the Public Health Enhancement Fund/South African National Department of Health. The views and opinions expressed are those of the authors and do not necessarily represent the official views of the SAMRC. We thank the mothers and their children for participating in the DCHS and the study staff, the clinical and administrative staff of the Western Cape Government Health Department at Paarl Hospital and at the clinics for support of the study. We also thank all research participants in the UK Biobank, BioBank Japan, PAGE study, UGR and Uganda GPC, and AGVP studies.

## Data and Code Availability

All data used in this study are publicly available. Data from the African Genome Variation Project was accessed by combining EGAD00010001045, EGAD00010001046, EGAD00010001047, EGAD00010001048, EGAD00010001049, EGAD00010001050, EGAD00010001051, EGAD00010001052, EGAD00010001053, EGAD00010001054, EGAD00010001055, EGAD00010001056, EGAD00010001057, and EGAD00010001058. The Drakenstein Child Health Study is committed to the principle of data sharing. De-identified data will be made available to requesting researchers as appropriate. Requests for collaborations to undertake data analysis are welcome. More information can be found on our website (http://www.paediatrics.uct.ac.za/scah/dclhs). Uganda GPC genetic data used in this paper were accessed through EGAD00010000965 and phenotype data was accessed via sftp from EGA (reference: DD_PK_050716 gwas_phenotypes_28Oct14.txt). We accessed data from the UK Biobank with application 31063. BioBank Japan summary statistics were accessed from http://jenger.riken.jp/en/result. GWAS summary statistics for the Population Architecture using Genomics and Epidemiology (PAGE) study were accessed through the NHGRI-EBI GWAS Catalog (https://www.ebi.ac.uk/gwas/downloads/summary-statistics).

All code used in analysis is available here: https://github.com/armartin/africa_prs.

