## Supplementary Figures 1-16, captions for Tables 1-7 for "Low generalizability of polygenic scores in African populations due to genetic and environmental diversity"

### Supplementary Information

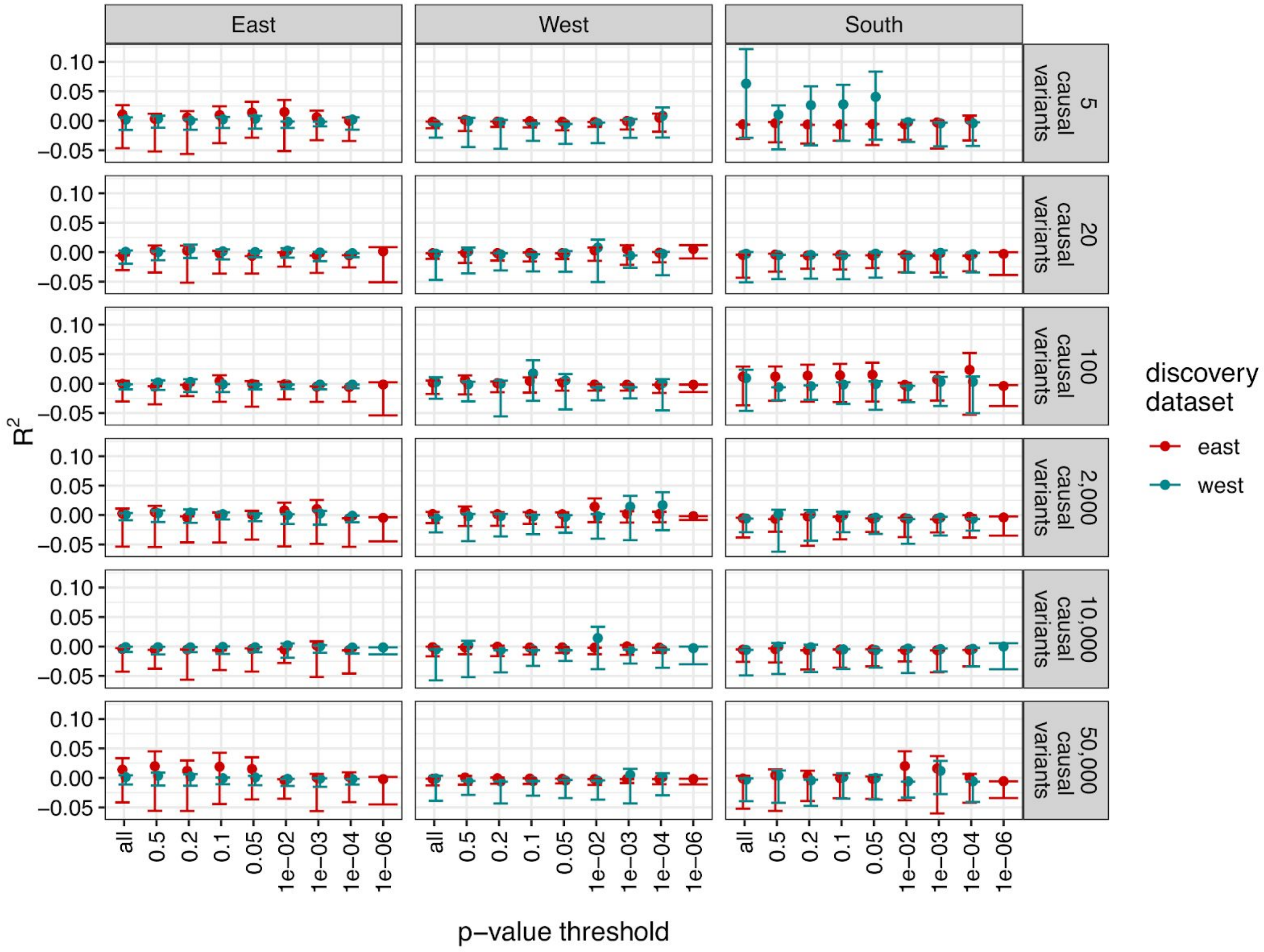


**Supplementary Figure 1 - Simulated polygenic score accuracies across regions of Africa using AGVP genotype data when $h^2$=0.1, similar to Figure 2.**

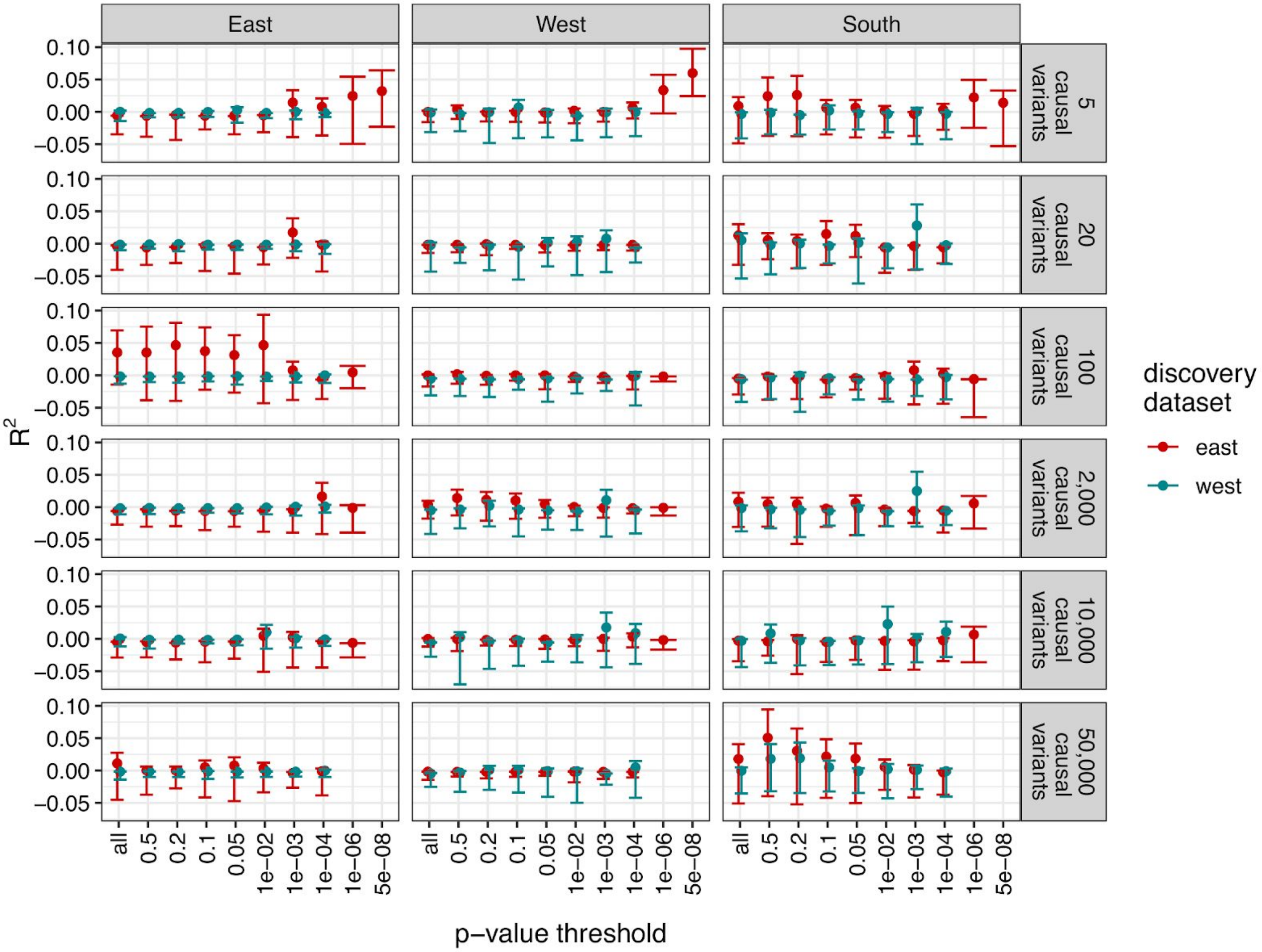


**Supplementary Figure 2 - Simulated polygenic score accuracies across regions of Africa using AGVP genotype data when $h^2$=0.2, similar to Figure 2.**

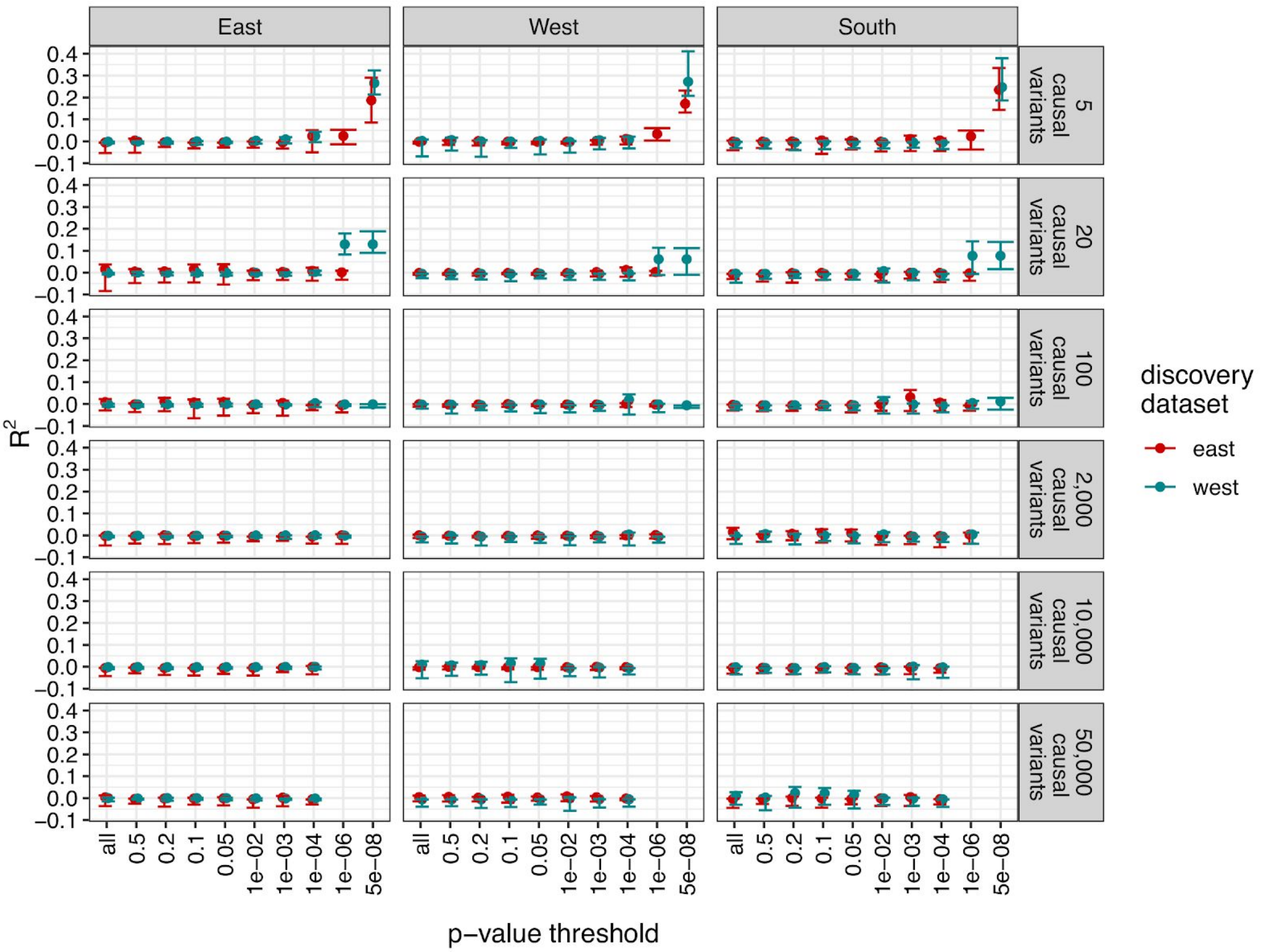


**Supplementary Figure 3 - Simulated polygenic score accuracies across regions of Africa using AGVP genotype data when $h^2$=0.4, similar to Figure 2.**

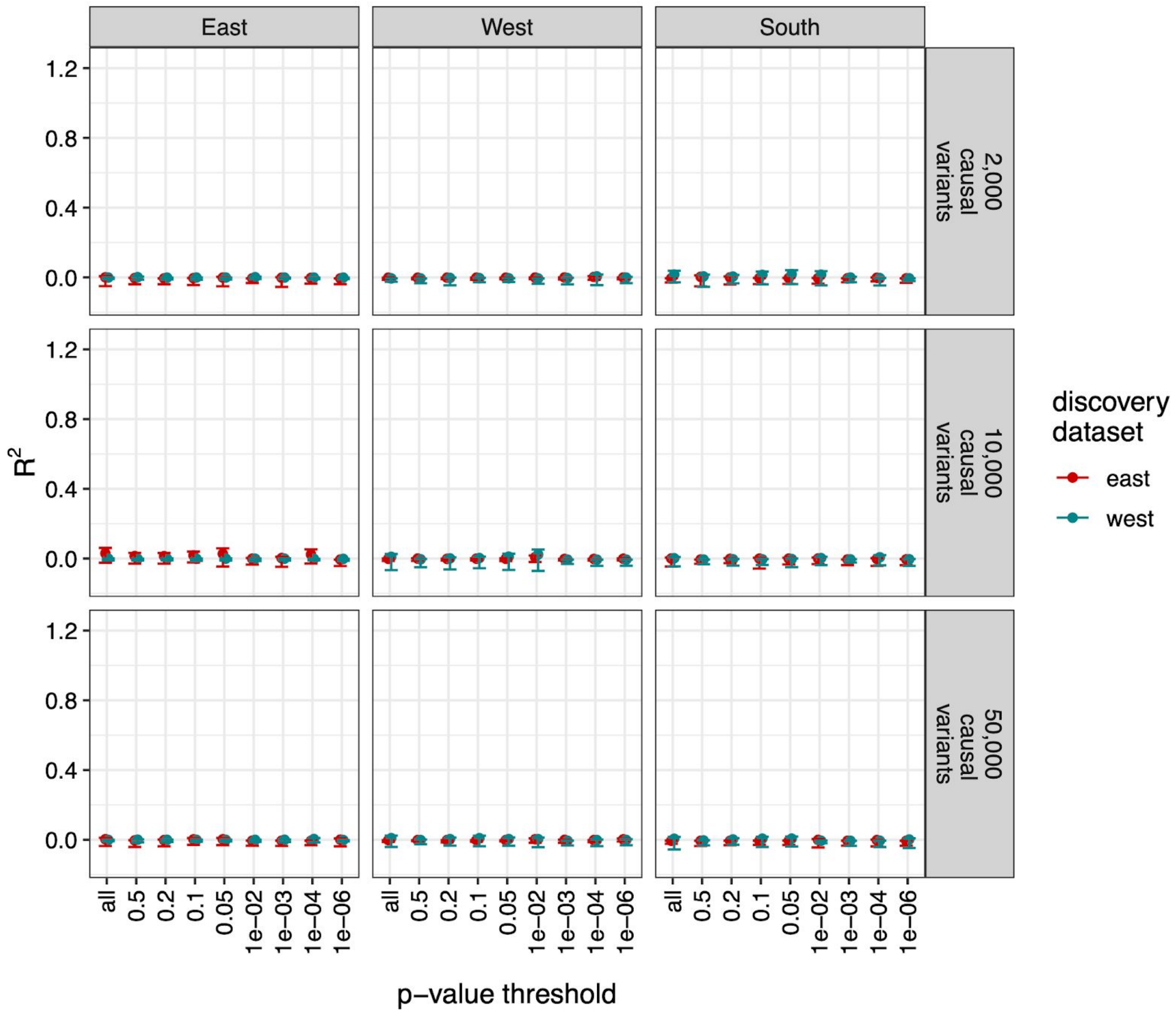


**Supplementary Figure 4 - Simulated polygenic score accuracies across regions of Africa using AGVP genotype data when $h^2$=0.8, similar to Figure 2.** For this degree of polygenicity, no variants were associated with the trait at a *p*-value less than 5e-08 hence this is not shown in the figure.

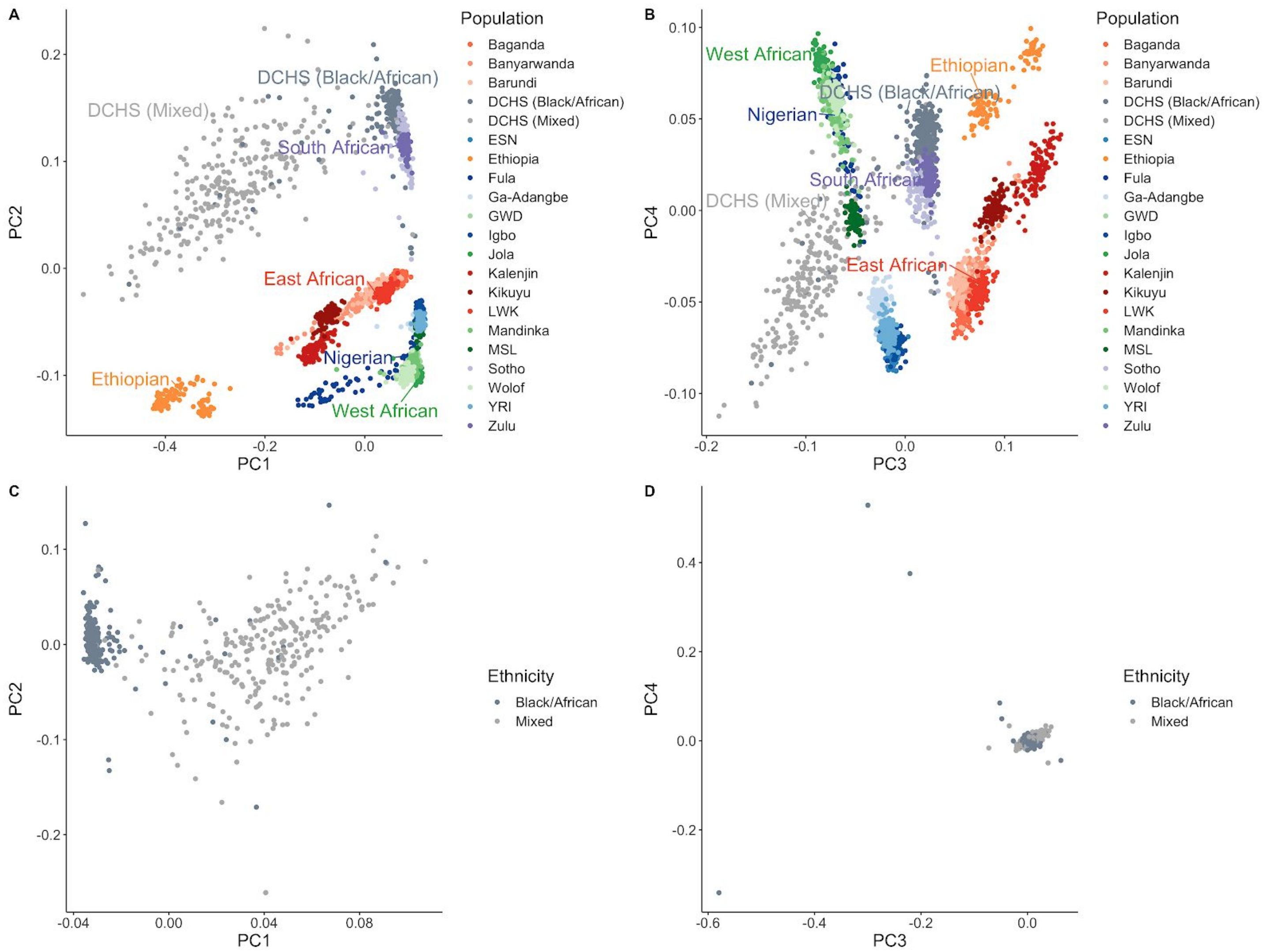


**Supplementary Figure 5 - Ancestry and ethnicity within DCHS and compared to African reference populations from AGVP and the 1000 Genomes Project.** A-B) PCs in DCHS Black/African and Mixed ethnicities compared to reference data. C-D) PCs within DCHS without reference data. A, C) PC1 and PC2. B, D) PC3 and PC4.

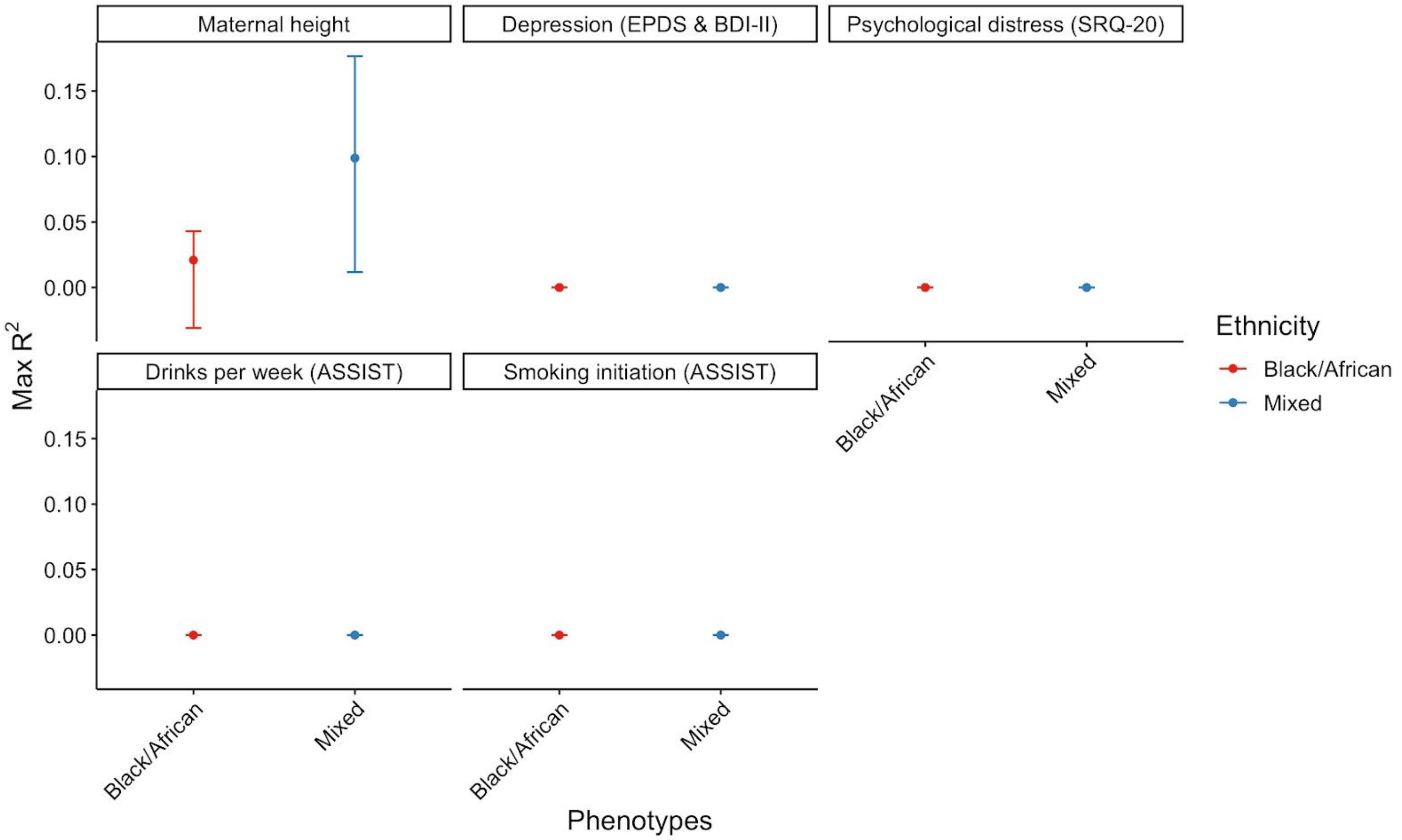


**Supplementary Figure 6 - PRS accuracy by ethnicity in the DCHS cohort for 6 different measured phenotypes.** Max $R^2$ shows specifically the PRS threshold that explains the most phenotypic variation. Only height is significantly predicted. Abbreviations as follows: EPDS = Edinburgh Postnatal Depression Scale, BDI-II = Beck Depression Inventory II, ASSIST = Alcohol, Smoking and Substance Involvement Screening Test.

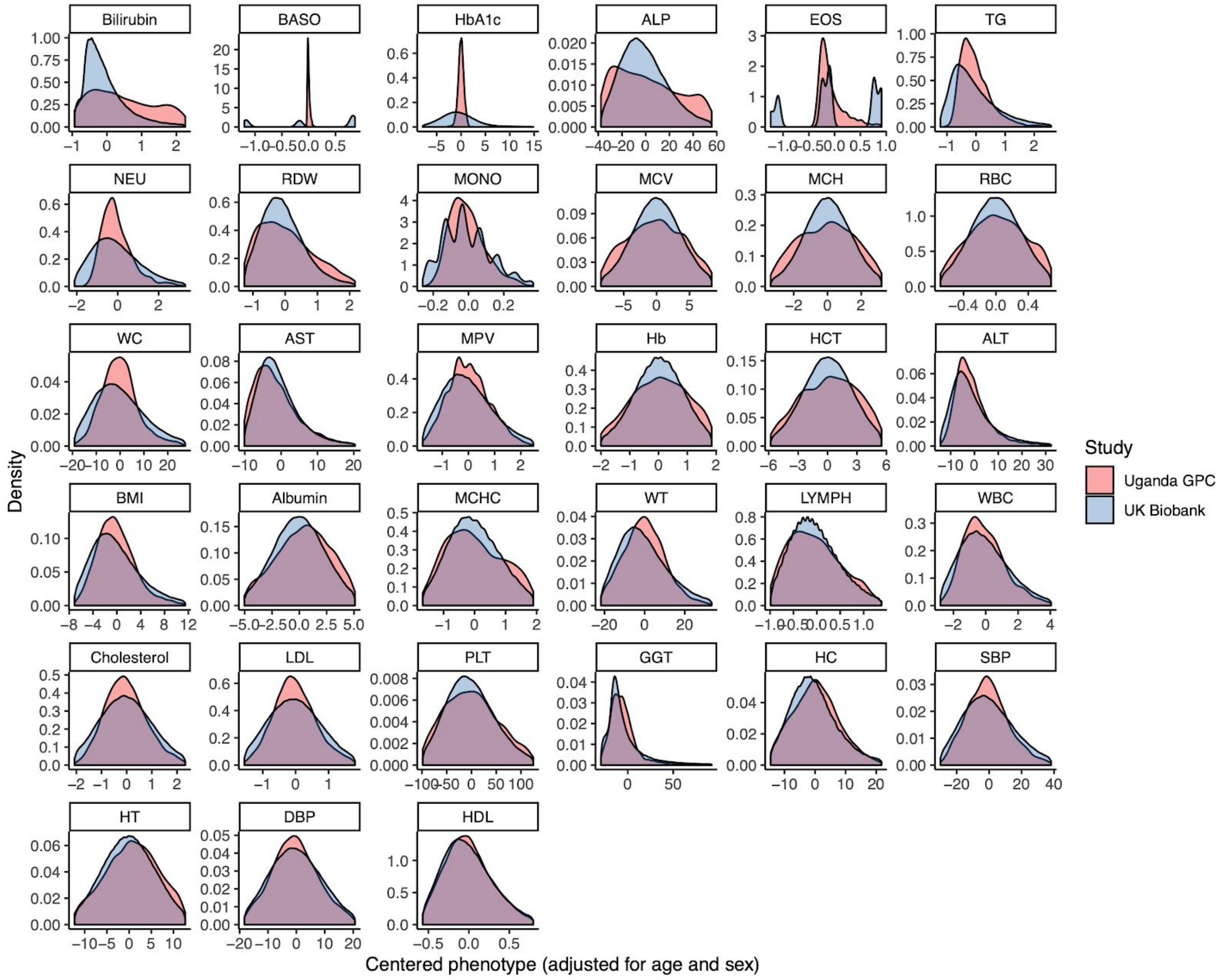


**Supplementary Figure 7 - Comparison of phenotypic distributions between the Uganda GPC and UK Biobank cohorts after mean centering phenotypes and regressing out effects of age and sex.** The middle 95th percentile of the data are shown for display purposes such that extreme outliers are removed. Phenotypes are ordered by distributional difference as estimated by the Kolmogorov-Smirnov test statistic.

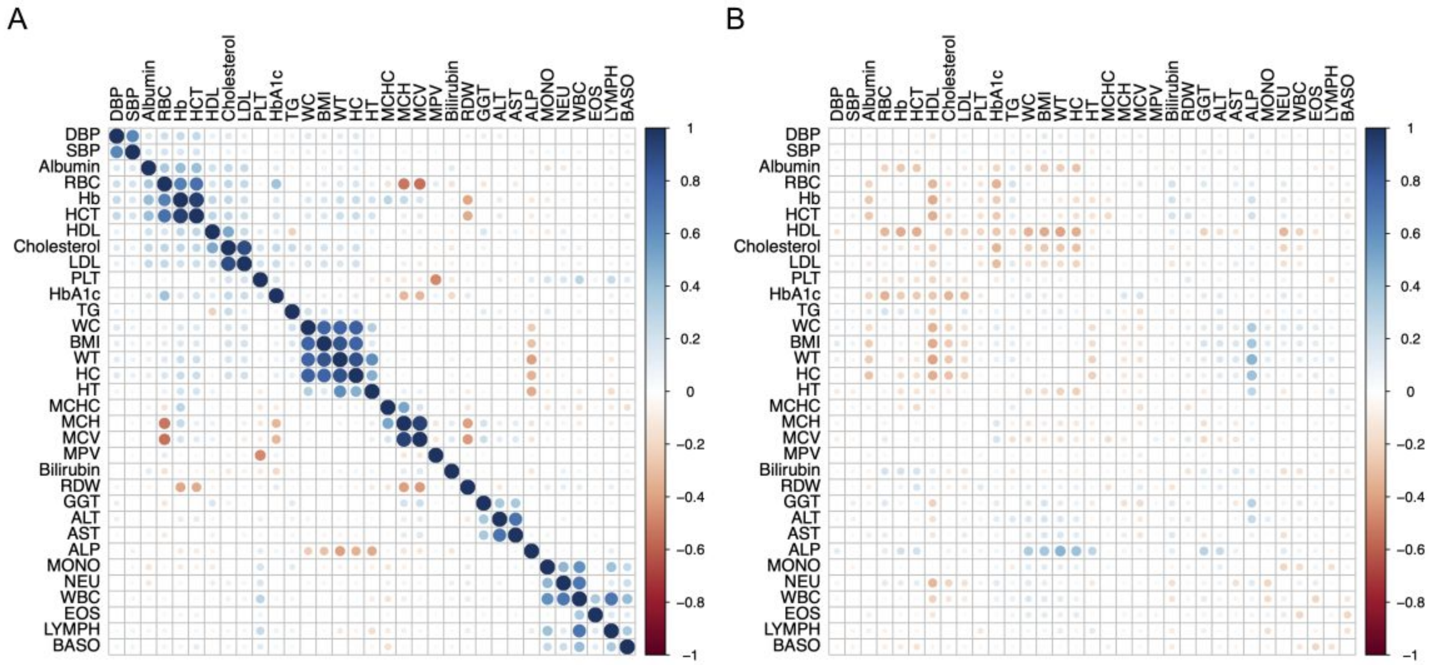


**Supplementary Figure 8 - Phenotype correlations among 34 quantitative traits measured in the Uganda GPC data versus UK Biobank data.** Analysis conducted as in Figure 4. A) Phenotypic correlation matrix when relatives are included in the Uganda GPC. B) Differences between phenotypic correlation matrices among unrelated individuals in the UK Biobank - Uganda GPC.

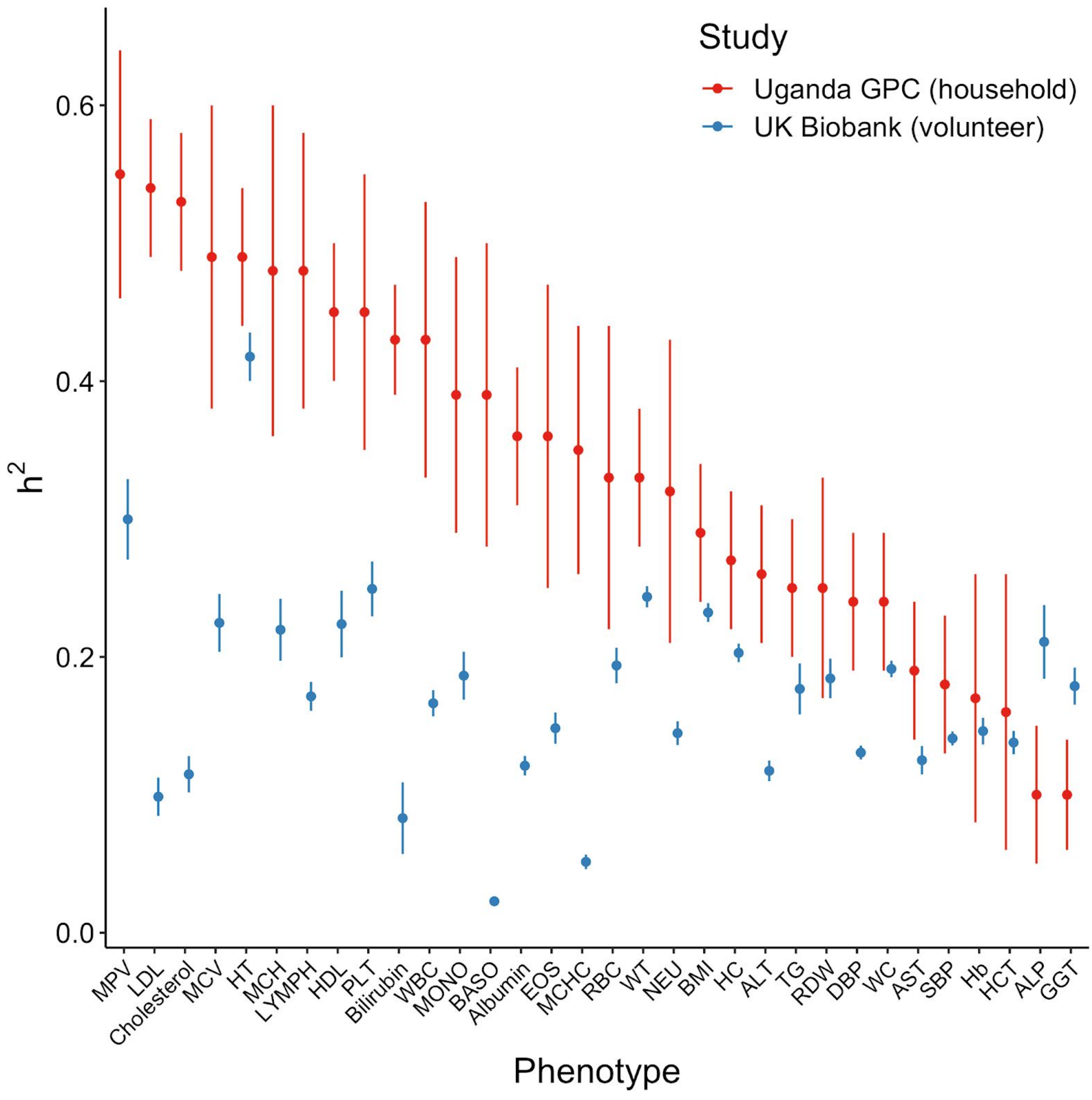


**Supplementary Figure 9 - Trait heritability comparison between UK Biobank and the Uganda GPC.** Note that we estimated heritability in the UK Biobank individuals using LD score regression from unrelated individuals, whereas a mixed model approach with multiple random effects were used previously in Uganda GPC due to the complex household and geographically diverse study design, as described previously(Gurdasani et al. 2019; Asiki et al. 2013; Heckerman et al. 2016). Given these differences, the higher but noisier heritability estimates in Uganda GPC are as expected.

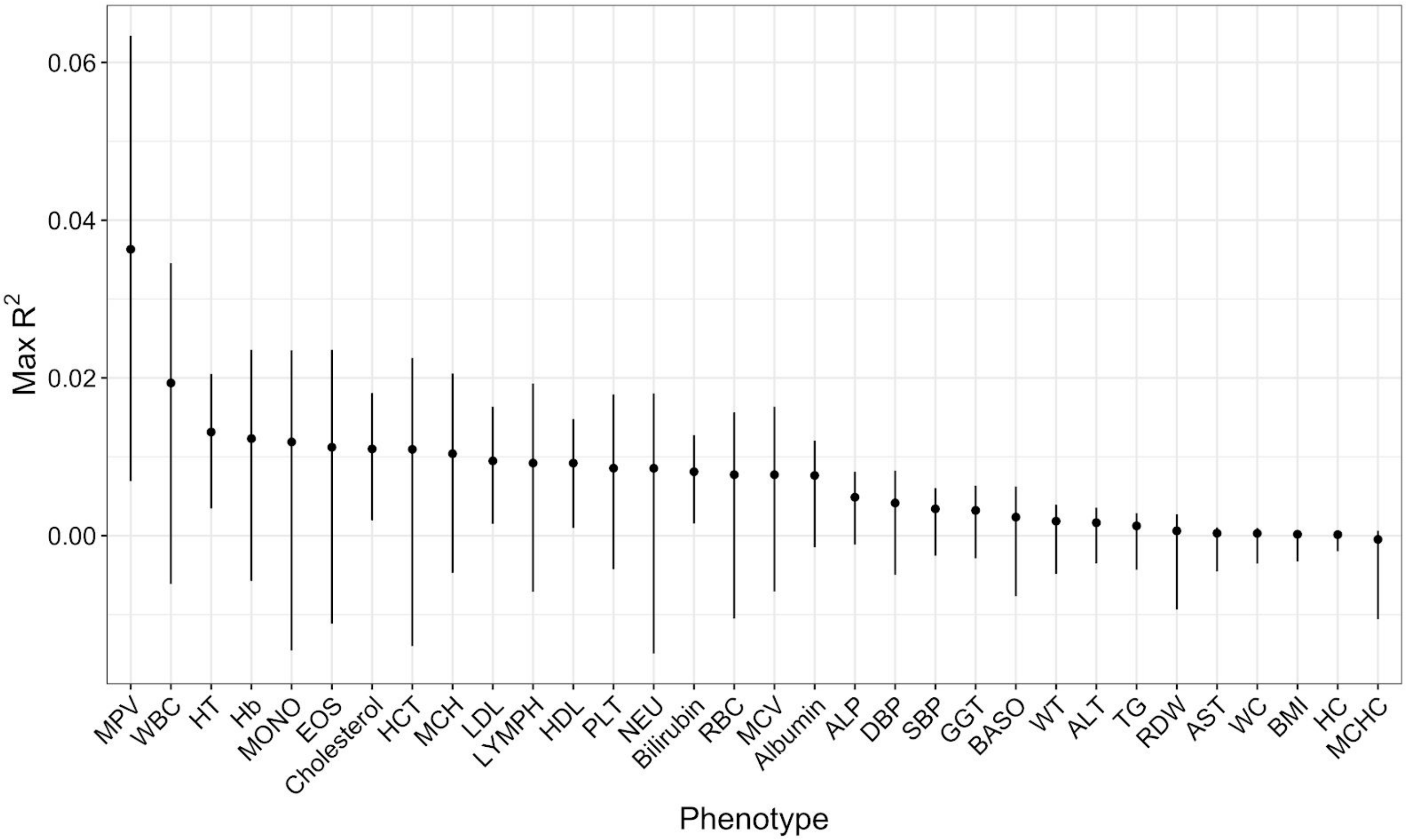


**Supplementary Figure 10 - PRS accuracy for 32 traits in unrelated Uganda APCDR individuals calculated using GWAS summary statistics from UK Biobank European ancestry individuals.** The PRS from the p-value threshold with the highest accuracy of 10 thresholds computed is shown.

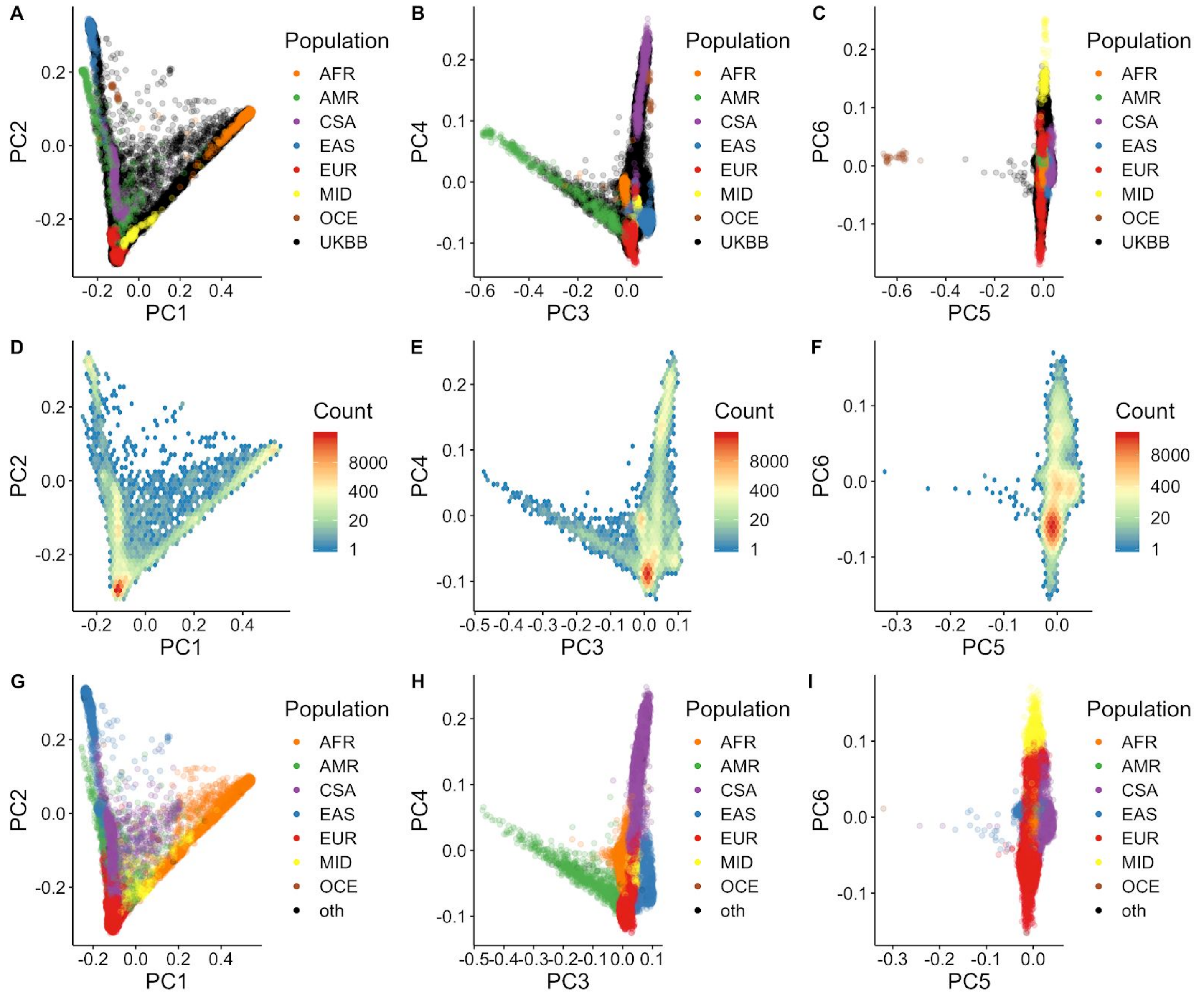


**Supplementary Figure 11 - Continental ancestries in the UK Biobank data with reference data from the 1000 Genomes Project and Human Genome Diversity Panel (HGDP).** A-C) Principal components analysis biplots (PCs 1-6) with loadings defined by reference data from 1000 Genomes and HGDP with colors corresponding to continental ancestry meta-data from these projects. UK Biobank data (UKBB) is projected into the same PC space and colored in black. D-F) Density of UK Biobank data (excluding reference panels) in PC1-6. G-I) Continental ancestry assignments in the UK Biobank using a random forest trained on meta-data from 1000 Genomes and HGDP (excluding reference panels). “Oth” are individuals whose ancestry was not confidently assigned to any ancestry group.

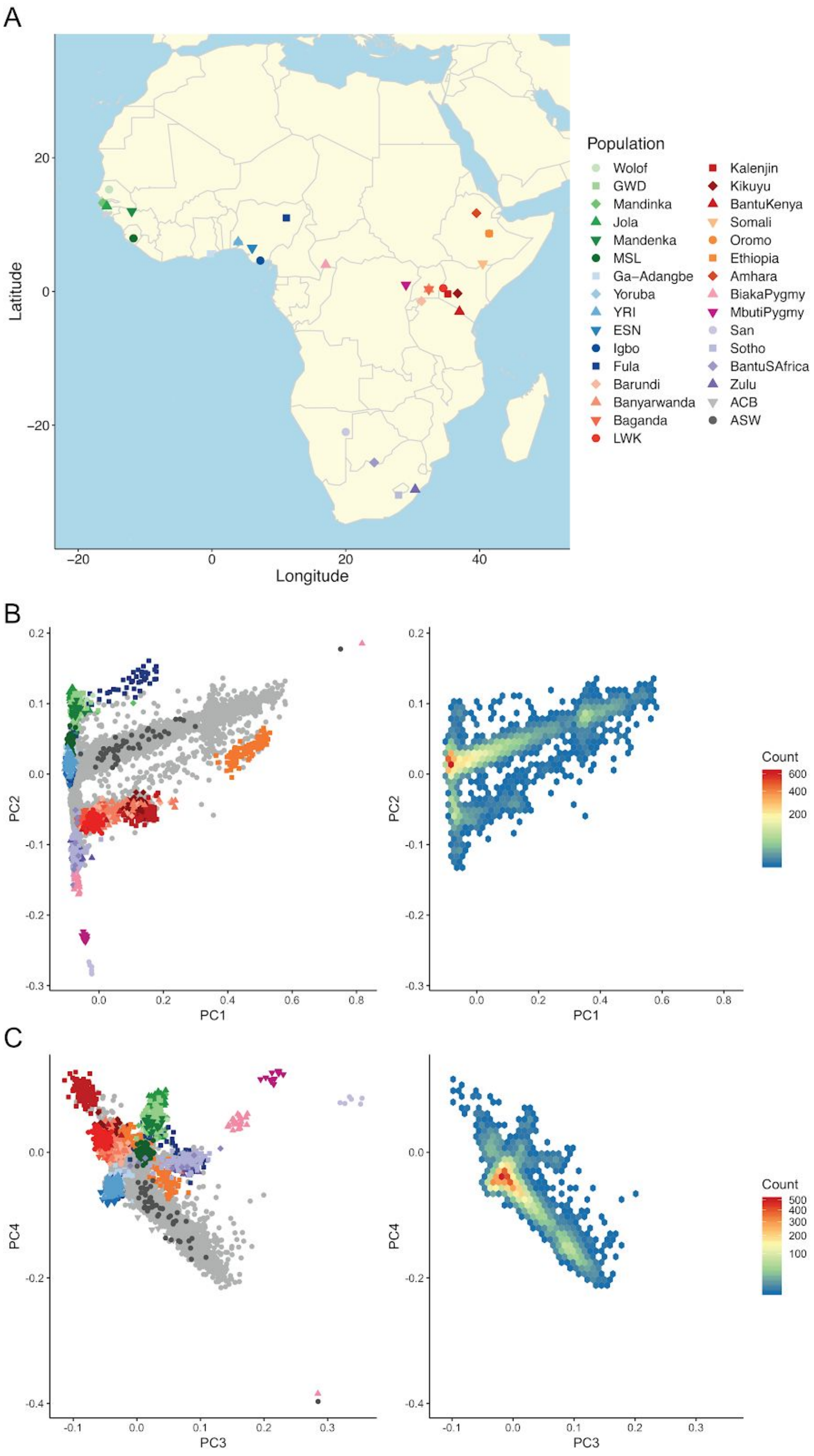


**Supplementary Figure 12 - African subcontinental ancestry in the UK Biobank**. A) Coordinates of African reference panel data from the African Genome Variation Project, 1000 Genomes Project, and Human Genome Diversity Panel. B) PC1-2 of UK Biobank and/or reference data. C) PC3-4 of UK Biobank and/or reference data.

B-C) Colors and shapes correspond to A), while grey points in the left plots are UK Biobank projected PC coordinates. Right plots show density of African ancestry UK Biobank individuals.

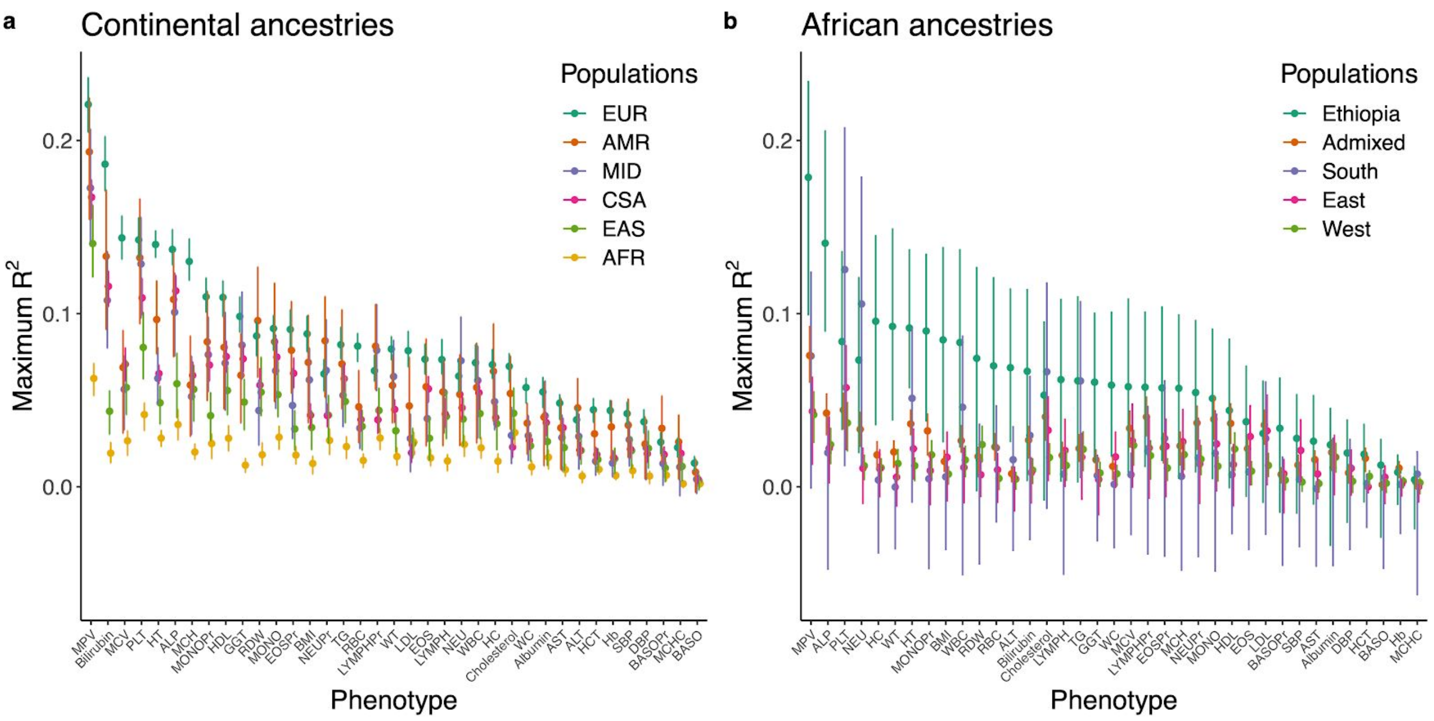


**Supplementary Figure 13 - Relative polygenic prediction accuracy across traits in the UK Biobank in target individuals**. Panels shown different A) continental ancestries, and B) African ancestries. All individuals in B) are from the “AFR” population in A). Only one PRS is shown from each trait, which includes the maximum $R^2$ threshold.

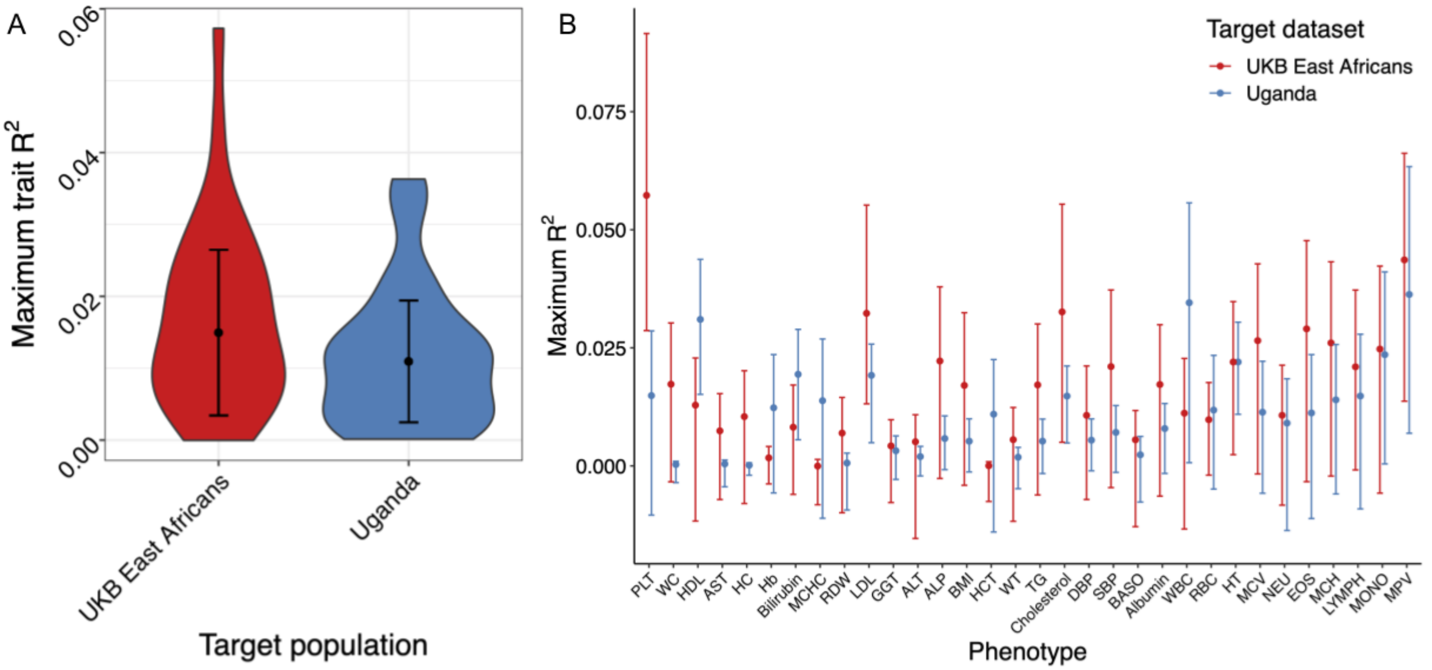


**Supplementary Figure 14 - Comparison of PRS accuracy for 32 traits across cohorts in individuals with East African ancestry.** Includes unrelated participants in UK Biobank (N=728 with East African ancestry) versus Uganda GPC (N=2,247), using the same summary statistics from UK Biobank European ancestry individuals (N=351,194). A) Summary of accuracy distributions across 32 traits. B) PRS accuracy in each trait. Traits are ordered by absolute difference in prediction accuracy among cohorts despite similar ancestries.

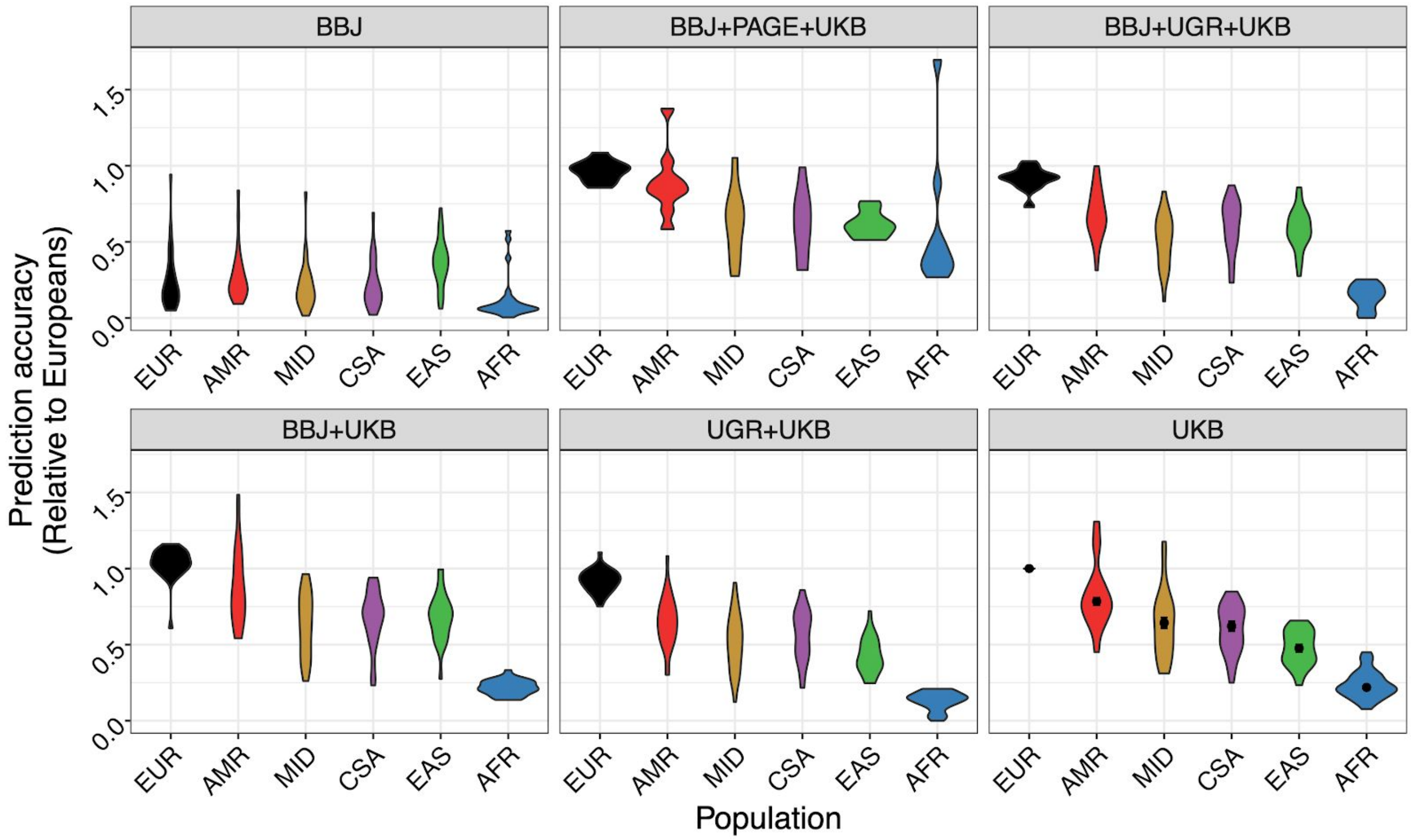


**Supplementary Figure 15 - Relative PRS accuracy using the same target individuals and varying discovery cohorts**. All relative comparisons are with respect to accuracy in withheld EUR when predicting with UKB GWAS summary statistics.

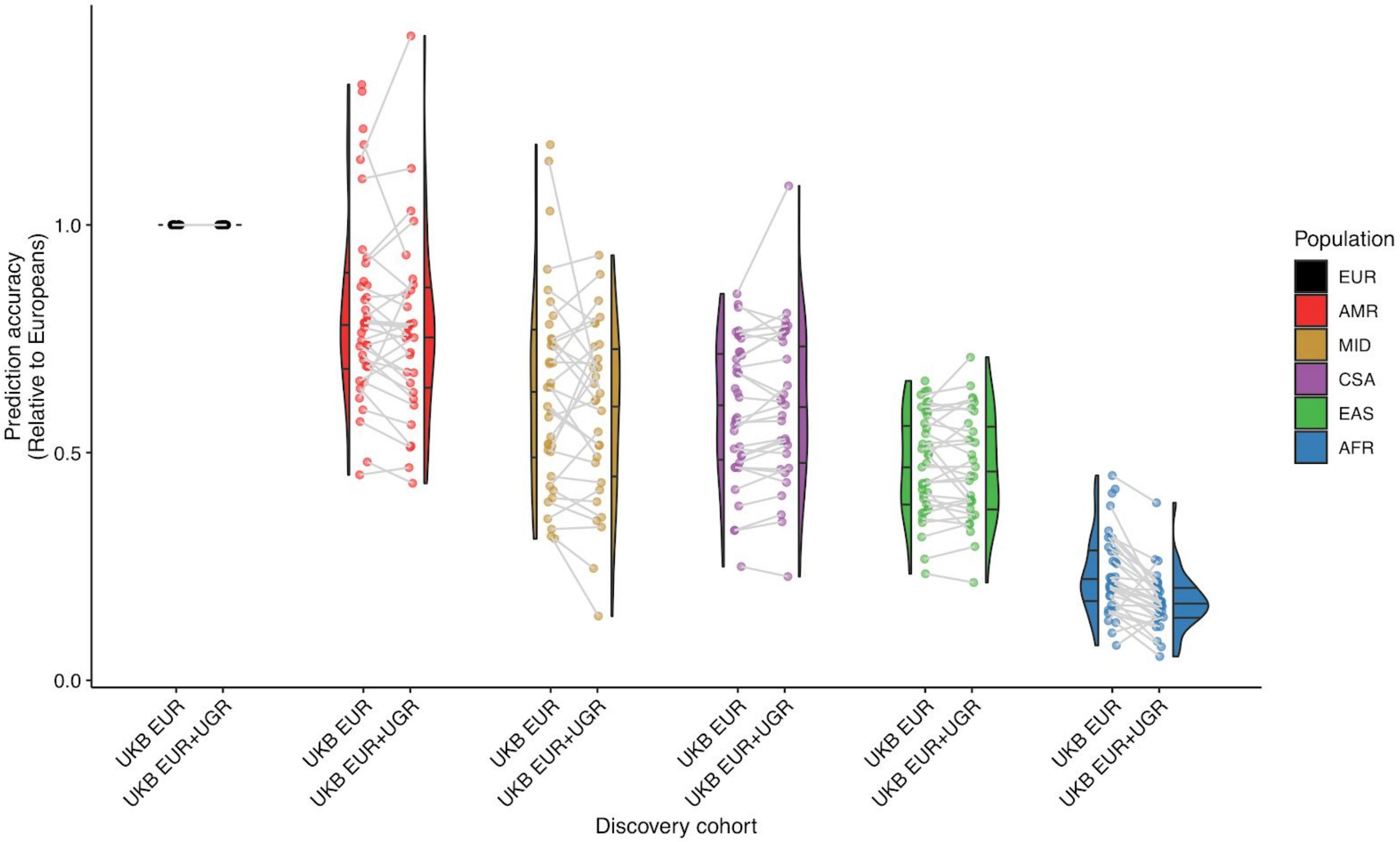


**Supplementary Figure 16 - Polygenic score accuracy in ancestrally diverse target populations from the UK Biobank versus meta-analysis combining UK Biobank with the Uganda Genome Resource (UGR) summary statistics.**

### Tables

**Supplementary Table 1 - Abbreviations used throughout this study.**

**Supplementary Table 2 - Overview of genetic and phenotypic data used throughout this study.**

**Supplementary Table 3 - DCHS phenotype descriptions and corresponding GWAS used for prediction.** Abbreviations as follows: EPDS = Edinburgh Postnatal Depression Scale, BDI = Beck Depression Index, ASSIST = Alcohol, Smoking and Substance Involvement Screening Test.

**Supplementary Table 4 - Uganda GPC phenotypes and corresponding GWAS discovery cohorts, including UKB, BBJ, and PAGE.** Kolmogorov-Smirnov (K-S) and F-tests were used to compare overall distributions and variances, respectively. CI=confidence interval. 95% upper and lower CI bounds for the F-test are shown. K-S test statistics provided the ordering of panels in **Figure S6**.

**Supplementary Table 5 - Counts of UK Biobank individuals assigned to each ancestry group at the continental level globally as well as regional levels within continental African ancestry.**

**Supplementary Table 6 - Meta-analysis discovery cohort summaries across phenotypes.** Number of individuals with genotype and phenotype data included in each GWAS discovery cohort analyzed. When constructing PRS from each meta-analysis combination for each phenotype, the fraction of individuals in the meta-analysis from each continental ancestry group was used to generate a weighted LD reference panel in the 1000 Genomes Project by matching with super population metadata. For this purpose, ancestry groups in GWAS were matched to 1000 Genomes super populations as follows: UKB was matched to EUR, BBJ was matched to EAS, UGR was matched to AFR, and PAGE was matched proportionally to AFR, AMR, and EAS by ancestry composition reported for each trait. For PAGE specifically, because the 1000 Genomes Project lacks Native American reference panels, Hispanic/Latino and Native American populations were summed and matched to 1000 Genomes AMR. The relatively small numbers of Native Hawaiian and Other were not matched to any group in 1000 Genomes for constructing LD reference panels.

**Supplementary Table 7 - Population-enriched variants with outsized impacts on genetic prediction accuracy for traits.** Lead SNPs come from associations that were genome-wide significant in both the meta-analysis and in UKB only ($p < 5e-8$). The top ten associated loci grouped by nearest gene were identified by modeling the linear relationship between -log10(p-values) in the meta-analysis versus UKB only, then filtering to only those variants above the 99% prediction interval (i.e. more significant in the meta-analysis than expected by a linear relationship), and then sorted by the difference in -log10(p-values) between the meta-analysis versus UKB alone.
